# Geometry aware graph attention networks to explain single-cell chromatin state and gene expression

**DOI:** 10.1101/2025.05.29.656611

**Authors:** Gabriele Malagoli, Patrick Hanel, Anna Danese, Guy Wolf, Maria Colomé-Tatché

**Affiliations:** Helmholtz Zentrum München, Institute of Computational Biology, Neuherberg, Bavaria, Germany; LMU Munich, Biomedical Center (BMC), Physiological Chemistry, Faculty of Medicine, Planegg-Martinsried, Bavaria, Germany; Université de Montréal, Department of Mathematics and Statistics, Montréal, Québec, Canada; Mila-Quebec Artificial Intelligence Institute, Montréal, Québec, Canada

## Abstract

High-throughput measurements that profile the transcriptome or the epigenome of single-cells are becoming a common way to study cell identity. These data are high dimensional, sparse and non linear. Here we present SEAGALL (Single-cell Explainable Geometry-Aware Graph Attention Learning pipeLine), a hypothesis free method to extract biologically relevant features from single-cell experiments based on geometry regularised autoencoders (GRAE) and explainable graph attention networks (GAT). We use a GRAE to embed the data into a latent space preserving the data geometry and we construct a cell-to-cell graph computing distances in the GRAE bottleneck. Exploiting the attention mechanism to dynamically learn the relevant edges, we use GATs to classify the cells and we explain the predictions of the model with XAI methods to unravel the features which are driving cell identity beyond marker genes. We apply our method to data sets from scRNA-seq, scATAC-seq and scChIP-seq experiments. SEAGALL can extract cell type specific and stable signatures which not only differ from the ones found in classical linear approaches but are less biassed by coverage and high expression.

## Introduction

Single-cell sequencing technologies have provided a breakthrough in molecular biology by allowing the measurement of transcriptomic and epigenomic profiles at the scale of single-cell with high resolution. Many reproducible and ready-to-apply kits have become common and affordable, leading to a great increase of interest in this field. For instance, single-cell RNA sequencing (scRNA-seq), also known as gene expression (GEX), or single-cell Assay for Transposase-Accessible Chromatin using sequencing (scATAC-seq)^1,2^ can be performed with the readily available kits commercialized by 10X Genomics by applying their well described protocols. A new step towards the understanding of molecular biology is the possibility to do multi-omics single-cell sequencing, which allows the measurement of more than one omics modality at the same time for the same single-cell. Among others, the 10X Genomics Multiome Platform, which quantifies the chromatin openness and the transcriptome of single nuclei, also allows to perform such measurements.

The standard analysis of single-cell data involves low-dimensional embedding, followed by cell clustering and identification of cell types^3^. The common assumption is known as the “manifold hypothesis”^4^: high dimensional data lie along a latent and unknown manifold with a smaller dimension than the observed space. In single-cell biology, we measure tens of thousands of variables, such as genes (for gene expression measurements) or genomic loci (for epigenomic measurements). These features cannot take any possible value, but rather they vary within well defined ranges given by biological constraints, like gene regulatory networks^5^. These constraints define the underpinning manifold whose exact equations are unknown. A single-cell experiment can be seen as a method to sample (cells) from this manifold. From the distances between cells we can create a graph that resembles the manifold as accurately as possible (Fig.1A).

**Figure 1:**
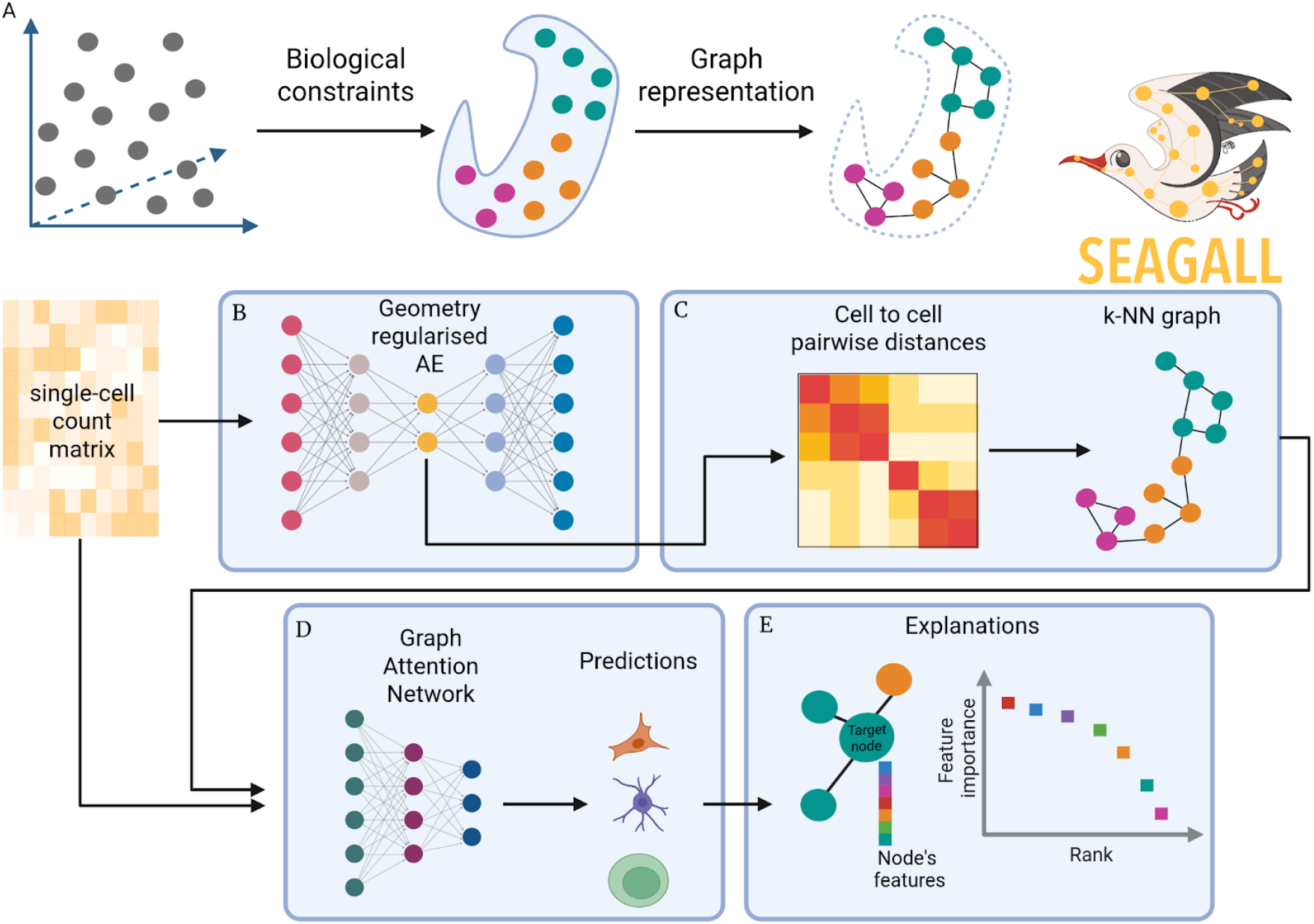
**A** without any constraint, the measurement of molecular features can take any arbitrary value (left). In reality, the gene regulation network imposes constraints which define the cell type and the possible values of the variables, defining a manifold on which the data live (center). As a proxy for the manifold it is possible to use a cell-to-cell graph, defined by pairwise distance between cells (right). **B** - **E** the SEAGALL model. The initial count matrix is reduced with a GRAE to preserve data geometry, i.e. both local and global structure of the data (**B**), within the latent space we compute pairwise distance between cells to build a cell-cell graph, (**C**). The graph and the count matrix are subsequently the input to a GAT classifier, whose predictions (**D**) will be explained in order to identify the most relevant features for each cell type (**E**).

Many tools exist to perform single-cell analysis^3^, yet they show several limitations. First of all, they are in general omic-specific^6–11^, relying on omic-specific assumptions, and forcing the users to choose a different tool for each omic. Moreover, most standard tools compute distances in a low dimensional linear space such as Principal Components (pca) or Independent Components (ICA)^6,8,9,12–14^. These linear assumptions lead to the loss of the intrinsic nonlinearity present in biological data sets, and prevent the discovery of complex insights between features and cells. Due to these shortcomings, autoencoders (AE)^15^ have recently become very popular because of their ability to learn the input and embed it in a nonlinear fashion^10,11,16–18^. Indeed, their strongest characteristic is the ability to take into account nonlinear dependencies within the data sets without making strong data assumptions. Yet, autoencoders often fail to represent the intrinsic data structure^19^, such as topology or geometry. To better fit the single-cell data, omic-specific AE have been developed; they assume a probability distribution from which the data are sampled and they use variational AE to embed the data sets^10,11,16,18^. As a consequence, a specific AE is needed for each modality, which grows in number day by day.

Finally, another important aspect of single-cell data analysis is the identification of features defining cell identity. The standard method to investigate what are the important features, such as genes or peaks, for each group of cells is based on differential analysis (DA). It consists in computing the distributions of the features in the different treatments, conditions or cell types to then quantify the difference between these distributions, giving as a result a list of features ranked by the most different to least ones. This approach will output features which are different between two groups of cells, but there is no guarantee that they are also relevant and important for each group specifically.

To address these limitations, we developed SEAGALL (Single-cell ExplAinable Geometry-Aware Graph Attention Learning pipLine), a deep learning method based on manifold learning and explainable AI for downstream analysis of single-cell data sets. SEAGALL first learns a low-dimensional embedding of the cells based on a graph-regularised autoencoder (GRAE)^19^ (Fig.1B). This embedding preserves both local and global structure of the data without particular assumptions on the data generation process. Then the tool computes the cell-to-cell graph on that low-dimensional space (Fig.1C), which is used as input to a graph attention network (GAT)^20,21^ together with the count matrix defining feature vectors of the nodes. The GAT classifies the cells into different cell types or states (Fig.1D) and the final output of SEAGALL are the explanations of the model, i.e. the set of input features which are the most important for the label prediction^22^ (Fig.1E). We applied our new method to ten different single-cell data sets spanning three omics (sc-RNAseq, sc-ATACseq, sc-ChIPseq) showing that it is able to reconstruct and embed the data, explain the cell types beyond common marker genes and extracting stable and specific features important for the cells which are not seen by standard differential analysis.

## Results

### The SEAGALL model

In a single-cell sequencing experiment, a set of genomic variables are measured for every cell, for instance gene expression in scRNA-seq, or openness of genomic loci in scATAC-seq. These measurements can be represented as a count matrix, where the set of specific variables are quantified within each cell. For example, in the measurement of the expression of *F* genes in *F* cells, the output is a *N* × *F* count matrix. The same holds for sc-ATACseq, where *F* is the number of loci in the genome for which the openness is quantified, called “peaks”. The count matrix can be seen as a method to store the position of *F* cells in an *F*-dimensional space, commonly called point cloud. Without constraints on the expression of genes or to openness of chromatin in the cells, the point cloud may span a homogenous volume in space. Yet constraints do exist, imposed for example by gene regulatory networks; therefore the data do not occupy a homogenous volume, but rather live on a manifold^23^, whose equations are unknown. However, the manifold is high dimensional, making little sense to compute distances on it due to the curse of dimensionality. Hence, the first step of SEAGALL is learning a low dimensional representation of the data that conserves the intrinsic geometry of the manifold, exploiting the recent development of geometry regularised autoencoders (GRAE)^19^ (Methods) (Fig.1B). GRAE first applies a kernel method named PHATE^24^ to learn the geometry of the data and uses it to regularise the structure of its latent space. Within the latent space of the GRAE, it is now possible to compute reliable pairwise distances between cells in order to create a cell-to-cell k-NN graph (Fig.1C). In the next step of SEAGALL, the cell-to-cell graph is used as input to a graph attention network^21^ (GAT) (Fig.1D), a graph neural network^25^ (GNNs) with an attention mechanism on the edges. The GAT is applied to classify the cells into multiple cell types, which are known based on previous cell type annotation. In this scenario the classification of a cell depends on its neighbourhood, via the joint embedding of features vectors, if the cell has degree (see Methods). The attention mechanism is important to dynamically learn the relevance of each edge: spurious edges will be ignored so the model can focus on the important ones. Finally, an explainable artificial intelligence method (GNNExplainer)^22^ (Fig.1E) is applied to explain the predictions: each cell is classified into a cell type and we can extract the most relevant features that the model has used to predict the label. This last step is critical to understand what are the most relevant genomic regions or genes that define cell types beyond the common markers.

### Model benchmarking

We carried out a breakdown of SEAGALL, testing each of its main parts: the embedding method (GRAE), the classifier (GAT) and the explainer (GNNExplainer). We used six count matrices for benchmarking the embedding method and the classifier, two from scRNA-seq and four from scATAC-seq (SuppTable1 and SuppTable2 for the cell type composition and dimensions). They are two multimodal data sets for which the scRNA-seq and the scATAC-seq were treated separately (human brain and human PBMC), the scATAC-seq part of a multimodal data set of mouse embryonic brain, and a scATAC-seq data set of kidney^26^ (see Methods for the count matrix construction and processing). We tested the ability of six embedding methods (see next paragraph) to recover the original data after adding artificial dropout and the quality of the cell-to-cell graph computed in the different latent spaces. To quantify the latter feature, we measure the homogeneity of the cell-to-cell graph in terms of cell type composition of the neighbourhood and the performance of a GNN classifier varying the input graph. We then tested the GAT and also a Graph Convolutional Network (GCN) architecture, computing F1 score accuracy, precision and recall of the classifiers. Last, we measured the stability and the specificity of the GNNExplainer. We also measured the classification and explanations performances of the final model on a scChIP-seq dataset of breast cancer (SuppTable1, SuppTable2), in which H3K27me3 was measured at the single-cell level^27^.

### Geometrical regularised autoencoders best recover corrupted data and capture biological structure

We benchmarked five different AEs architectures. We tested the GRAE together with a topological autoencoder (TopoAE)^28^, PeakVI^11^, scVI^10^, a standard variational autoencoder (VAE) and linear pca. The TopoAE was included to compare the GRAE to an AE with a similar rationale behind: while the GRAE regularises the loss function considering that the geometry of the data should be preserved in the latent space, the TopoAE preserves the topology of the input space by applying persistent homology. Geometry is a more specific and local property than topology; however neither GRAE nor TopoAE make assumptions about the data sets, which makes them applicable to, in principle, any kind of biological data. The VAE is taken as a baseline model of autoencoder, to compare sophisticated methods to a simpler one. PeakVI is a state-of-the-art scATAC-seq specific AE, tailored for this data type so it can only be applied to it. scVI is a single-cell AE which represents the state of the art to embed scRNA-seq data, it therefore can only be applied to this data modality. pca is included in the benchmarking to test how different a linear dimensionality reduction method performs. We quantified which AE architecture is better to use for dimensionality reduction of single-cell data to then construct the cell-to-cell graph.

To measure the AEs ability to retrieve corrupted data, we applied a variable dropout between 0% and 50% in regular incremental steps of 10% to the six RNA and ATAC count matrices and trained each AE on the faulty data. We repeated the experiment ten times to ensure that each time the dropout will affect different features and to have a statistically reasonable sample size. After the training we measured the mean squared error between the decoded matrices of each AE and the original, non-corrupted, matrices (Fig.2A, Methods). GRAE outperforms all the methods achieving the minimum MSE at every dropout level (Fig.2B-G), except for the highest dropout on the peaks of the PBMC data. In particular, the geometry regularised AE is better than the two -omic specific AE scVI and PeakVI.

**Figure 2:**
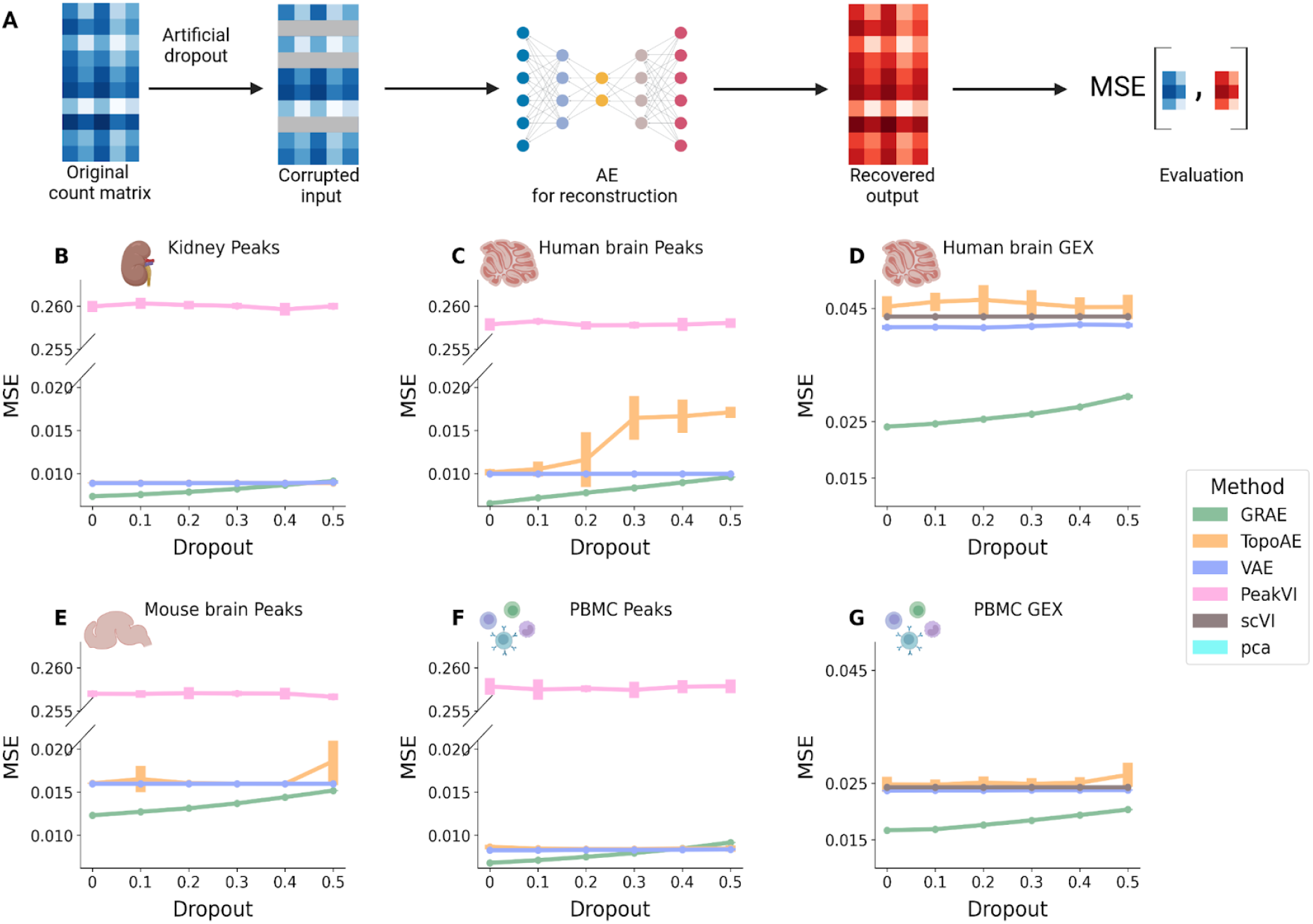
**A** schematic of the benchmarking process: data is corrupted with artificial dropout and the AEs reconstruct them. We evaluated the performance computing the MSE between the recovered output and the original count matrix. **B-G** average MSE at different levels of artificial dropout for each AE. Each point is the average of ten runs and the height of the error bar represents three times the uncertainty on the mean.

Then we measured the homogeneity of the k-NN graphs built from the AEs latent spaces. We assume that a good latent space leads to a k-NN graph where neighbours of a node belong to the same cell type. The more homogeneous the neighbourhood, the more the AE is able to locate close to each other in the latent space cells sharing the same biological functions. We trained each AE and we computed the k-NN (k=15) graphs from their latent spaces. Then, we measured for each cell how many cell types are found in its neighbourhood. We normalised this value by the product of the number of the neighbours and the number of cell types within the data set. We call this score “heterogeneity”. The value reported for each AE and each count matrix is then the average “homogeneity”, which is computed as one minus the average heterogeneity for each neighbourhood (Fig.3A, Methods).

**Figure 3:**
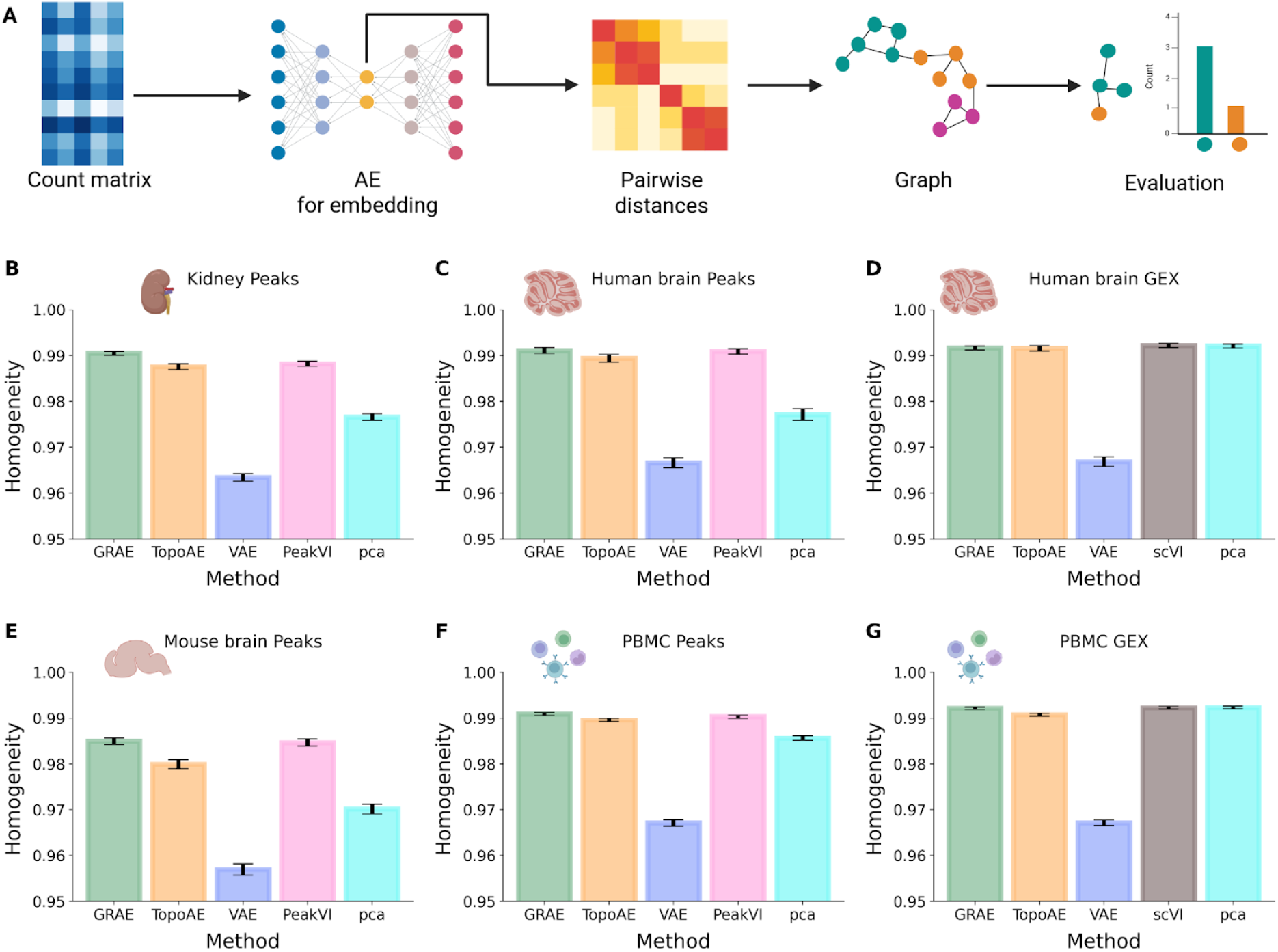
**A** schematic of the benchmarking process: each count matrix is embedded using the different AEs, the cell-to-cell graph is computed from the latent space and its homogeneity is used to evaluate the performance of the AE. **B-G** homogeneity of the k-NN using the different embedding methods. The height of the bar represents the average homogeneity across runs and the error bars spread is three times the uncertainty on the mean.

For the ATAC data sets, GRAE outperforms all the methods on the kidney data set and on the ATAC part of the PBMC data set (Fig.3B,F); on the other two ATAC count matrices, PeakVI and GRAE achieve the same homogeneity (Fig.3C,E). The homogeneity of the graphs computed from GEX count matrices is similar between GRAE, scVI and pca (Fig.3D,G, SuppTable4).

In conclusion, GRAE, which applies a geometrical regularisation to the loss function, outperforms all other methods when it comes to reconstructing the initial input, even with the addition of noise in the data. It also performs either better or equal to the other methods in recovering the cell type composition in the latent space.

### Geometry aware graph attention networks achieve best classification performances

Last, we tested together the performances of different embedding strategies and GNN classifiers. We used the aforementioned AEs and pca as dimensionality reduction (DR) methods, combined with two types of GNN: GAT^20^ and GCN^29^. Each combination of the DR method and GNN with the explainer was run fifty times, changing the initial seed, to statistically test the stability of the results. For each combination and each run we applied six metrics: four metrics for the classification performance (accuracy, F1 score, precision and recall) and two metrics to measure the quality of the explanations (specificity and stability) (Fig.4A). Specificity quantifies how specific the explanations are for each cell type. It is defined as one minus the average overlap between the most important features for each cell type with the most important for each other cell type. Stability measures how much the explanations change by running a new instance of the same classifier and explainer for the same count matrix. It is computed as the average intersection between the extracted features of a cell type across different repetitions of the same classification and explanations (Methods). We did not measure any statistical differences between the performances of GAT and GCN in terms of F1, accuracy, precision, recall, specificity and stability (SuppFig1, SuppTable4). Meanwhile, the GRAE graph outperformed that of PeakVI, TopoAE, VAE and pca in terms of F1 score, accuracy, precision and recall (Fig.4B,C,D,E, SuppTable5). The scVI graph outperformed the one from GRAE on most of the metrics; however scVI is sc-RNAseq specific, preventing the possibility to apply it to every single-cell data The specificity and the stability of the explanation are very high with all the embedding methods, with GRAE either leading or being the second in the rank (Fig4F, G, SuppTable5). We tested the final combination of GRAE and GAT on the scChIP-seq data set, and showed a very high performance also for that data type in terms of accuracy, F1, precision and recall, as well as specificity and stability of the discovered features (SuppFig2).

**Figure 4:**
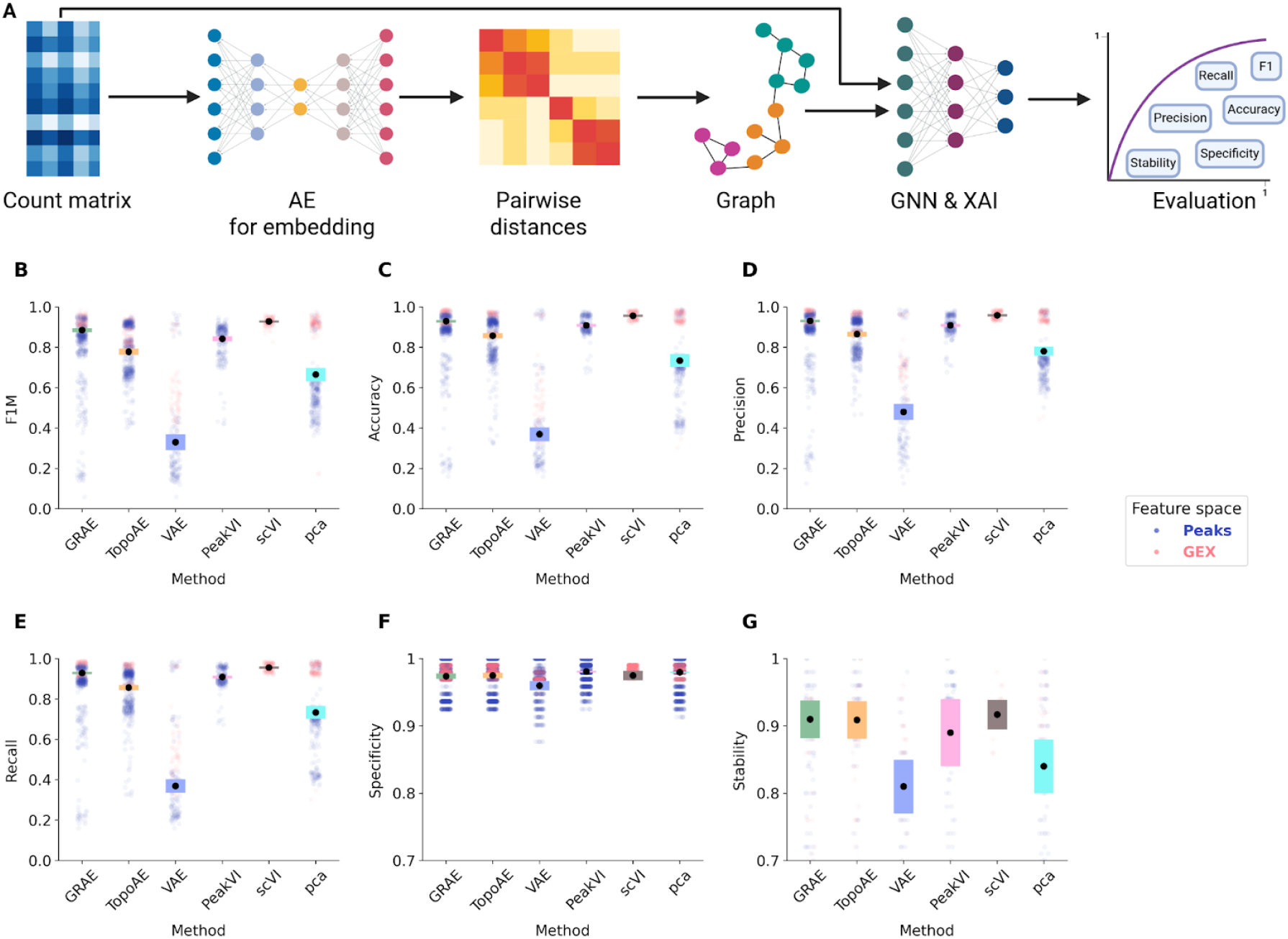
**A** schematic of the benchmarking process: After computing the k-NN graph, we trained a GNN classifier and applied the GNNExplainer; we finally computed the shown metrics to evaluate the models. **B** - **E** classification performances of the GAT classifier varying the embedding methods. Each black dot represents the mean across 50 runs and the height of the bars represent three times the standard deviation of the mean. **F** - **G** specificity and stability of the explanations. The vertical axes start from 0.7 for visualisation purposes. In all panels, each pale plot represents the point contributing to the mean, colored by data modality. PeakVI is only run on peak data sets and scVI is only run on GEX data sets; all other methods are run on all data sets.

In conclusion, these results indicate that the GRAE combined with a GNN classifier outperforms all other methods except of scVI for classifying cells into cell types and have the most stable and specific explanations.

### SEAGALL retrieves stable, specific and unbiased features

Given these results, the final SEAGALL model consists of the GRAE to embed the data and build the graph, and the GAT to classify the cells (Fig.5A). However, the last and crucial step is the explanation of the predictions. This point is crucial since it moves the focus from prediction performances to model interpretability, making the tool translational and useful for providing new biological insights. Once the GAT is trained on the geometry aware graph, SEAGALL investigates what are the features which are driving the predictions of the model, assuming that these features are the most relevant for the cell type and can define it beyond most common marker genes. To address this point, it applies a mask-based graph neural network explainer, known as GNNExplainer^22^. Given a node *v*_*i*_ the explainer finds the subgraph *G*_*s*_ and the subset of features *X_s_* = {*x*_*j*_|*v*_*j*_ ∈ *G*_*s*_} that maximises the probability of having the observed prediction 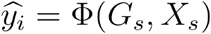 where Φ is the trained GNN. In other words, the explainer finds the subset of nodes features (and a subset of nodes links) that are most important to predict the nodes label. The importance is defined as the mutual information between the feature and the predictions (Methods). The distribution of the features importance drops fast with the rank, especially in GEX data. For both genes and peaks, at around the two hundredth feature, the importance drops one order of magnitude (Fig.5B, SuppFig3). Therefore, we suggest keeping a lower number of features for downstream analysis. For the single scChIP-seq data set, window features show a different behaviour: the maximum importance is significantly smaller than in peaks and GEX, and the importance decay is slower (Fig.5B). We speculate that because of the more noisy and sparse nature of the data each individual feature has a lower impact on the final prediction. However, the rate decay of the importance, computed as the absolute value of the derivative of the importance by the rank, shows that for all the three feature spaces the importance of the features does not change anymore after rank two hundred. Windows have a slower decay, again suggesting that each individual window has a lower impact on the results (Fig.5C). We picked fifty features for the downstream analysis, where we quantify the impact of technical biases on the DA and XAI. However, the influence of this threshold on the following results is very limited, within the range of 10-200 features (SuppFig4). We measured the stability and specificity of the XAI features (XAIFs). Stability is defined as the average overlap between the explanation of the same cell type running a new instance of the model and the explainer, therefore it measures the robustness of the method. Specificity is defined as the average overlap of explanations between cell types, quantifying the ability of the method to retrieve different features for different cell types. Our method is able to extract cell type specific (Fig.5D) and highly stable (Fig.5E) features in all the ten count matrices we tested. This means that it can consistently understand and explain a data set in order to suggest the key degrees of freedom which can be studied in dowsteam analysis and wet lab experiments, linking the deep learning method to real-world information. Notably, the XAIFs consistently differ from the differential features (DAFs) (Fig.5F), providing a potential for the discovery of novel data characteristics. This is due to the non-linearity and awareness of geometry of the model we propose, as replacing either GRAE with pca or GAT with standard NN leads to different explanations (SuppFig5). A direct comparison between XAIF and DAF shows that the former are less biased by high openness or expression and coverage (Fig.5G,H). This is particularly strong with peaks. Nevertheless, the lower biases of XAIFs are not traded off with a higher noise: the signal to noise (STN) ratio of XAIFs and DAFs is, on average, the same in all the feature spaces (Fig.5I).

**Figure 5:**
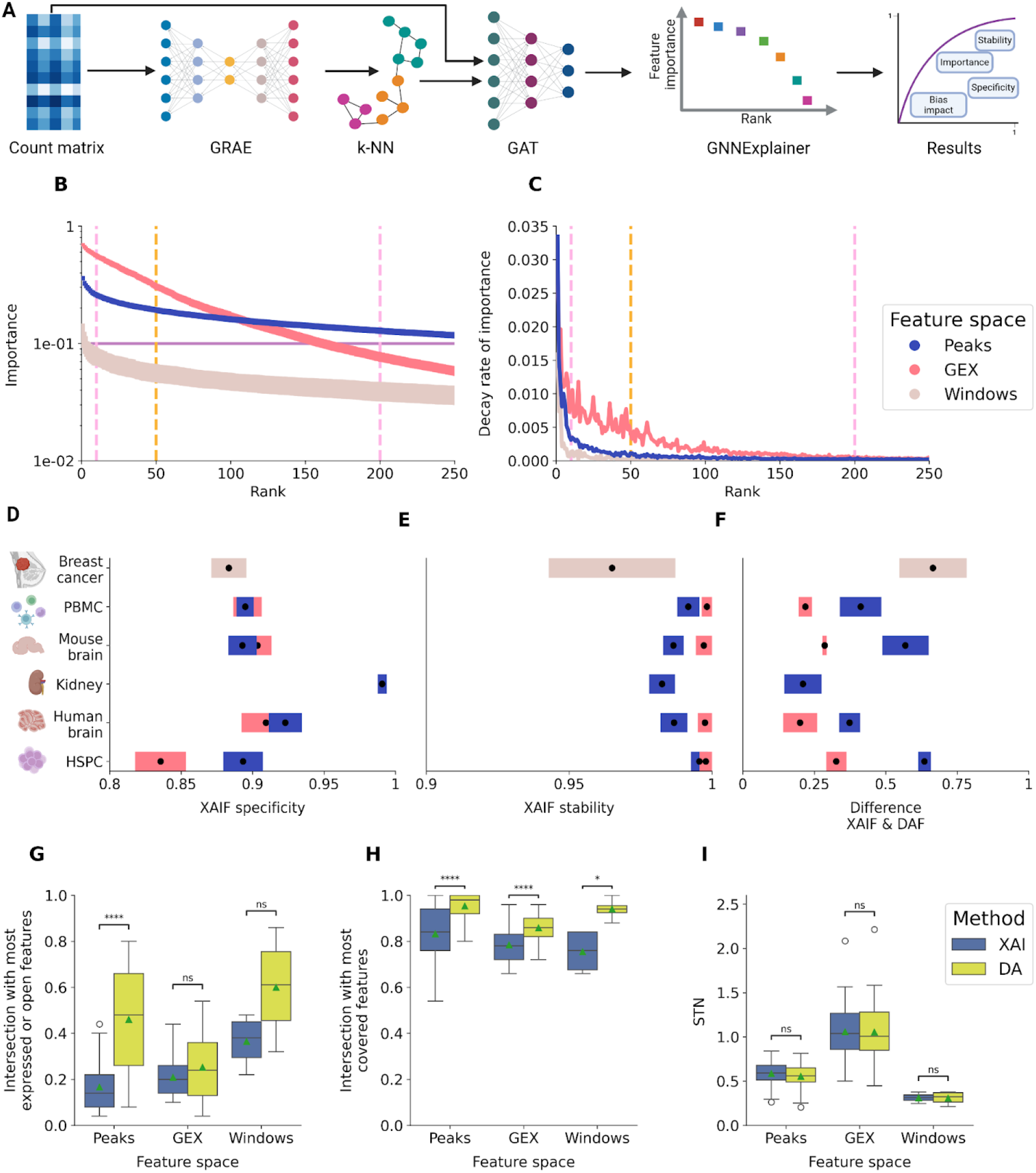
**A** final workflow of SEAGALL. **B** rank-importance distribution of the features according to the explainer. Average across data sets and cell type. Vertical dashed lines highlight the interval 10-200 features within the importance is stable. **C** Decay rate of the importance of the features, legend as in A. **D - F** specificity (left), stability (center) and difference between xai features (XAIF) and differential features (DAF) (right) of the explanations on the ten count matrices, spanning three different feature spaces and using 50 features for each label, legend as in A. **G** distribution of the overlap between XAIF and most expressed or open features (blue) and overlap between DAF and most expressed or open features (yellow). **H** distribution of the overlap between XAIF and most covered features (blue) and overlap between DAF and most covered features (yellow). **I** distribution of the signal-to-noise (STN) ratio of XAIF and DAF.

### SEAGALL identifies chromatin priming states and known cell type predictors

To study the biological significance of our results, we studied the features which were identified only by SEAGALL and not by differential analysis. For the scATAC-seq feature spaces, to link genomic loci to transcription factors (TFs), we run a motif analysis on the top important features, using HOMER^30^.

HOMER takes as input a set of genomic intervals and identifies enriched motifs, i.e. recurrent patterns of bases, and it checks whether these patterns match known motifs of TF binding. In the human brain data set we explored both the GEX and ATAC modalities. Taking the scATAC-seq XAIF and running motif analysis, we identified several brain specific motifs, which were not retrieved by motif analysis on the DA specific features (ExtendedTable1 for the complete motif results). In the astrocytes progenitors, SEAGALL could identify motifs belonging to the well known family of TFs SOX, such as SOX9 (Fig.6A), SOX17 (SuppFig6A) and SOX1 (SuppFig6B). SOX9 is known to be essential for the correct development of astrocytes^31^ and its promoter is activated to determine astrocyte differentiation^32^. Notably, the two dimensional embedding obtained with GRAE can well capture the differentiation process from astrocytes progenitors to astrocytes along its horizontal axis (Fig.6B). We measured the openness of all SOX9 TFBS (obtained from HOMER) and it turns out that they are already open in the astrocyte progenitors with a maximal openness in astrocytes (Fig.6C). On the other hand, the scRNA-seq modality shows that the expression of SOX9 is very limited in the progenitors but very high in the mature cells (Fig.6D). The other motif we retrieved is SOX17 (SuppFig6A), which is a TF known to be upregulated in astrocytes^33^. We discovered the openness of its TFBS as relevant for the identity of astrocyte progenitors; hence, we found a relevant TFBS openness in a progenitor population which is related to the expression of the TF in the direct next cellular state. For both SOX9 and SOX17 we therefore see the relevance of chromatin state priming the gene expression in progenitor cells, as suggested in^34^. Standard differential analysis could not highlight this dynamic behaviour. Focusing on GEX, SEAGALL ranked in the top fifty features of astrocytes the genes DNAH7 and EFEMP1, which were not identified with standard differential analysis. The former is known to be expressed in intermediate astrocytes^35^ and the latter is known to be expressed during synaptic development of astrocytes from iPSCs^36^. We correctly identified these genes during their positive gradient expression from the astrocytes progenitors to the astrocytes (Fig.6E,F). In addition, reprogrammed astrocytes have been shown to express a SOX1 positive state with neuronal stem cells characteristics^37^ and we identified its motifs (SuppFig6B) within the XAI features. Also in the human brain data set, only SEAGALL was able to obtain the motifs of JUNb, FOSL2 and FOS (SuppFig6C,D,E) as enriched among the discovered XAIF for microglia in the ATAC modality. These TFs are known lineage determining for microglia^38^. For GEX in microglia only our method highlighted two important genes, TLR2 and RIPK2 (SuppFig7B,C): the former modulates microglial activity^39^ and the latter plays an essential role in the microglia inflammatory response^40^. Last, in the brain cells annotated as inhibitory neurons, we found the motif of ASCL1 (SuppFig6F), which is known to specify and promote differentiation of GABAergic interneurons (i.e. inhibitory neurons)^41^.

**Figure 6:**
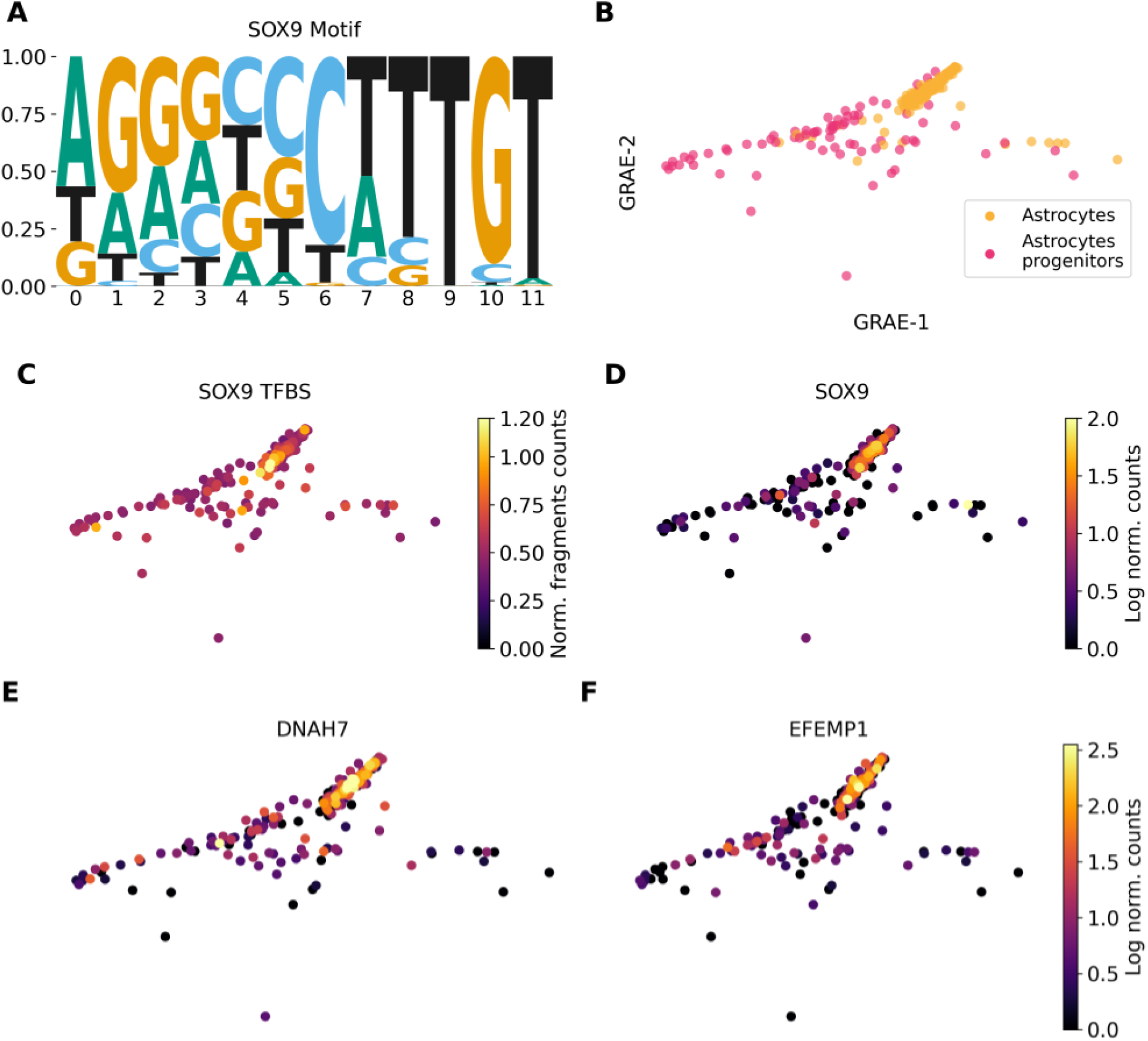
**A** SOX9 motif. **B** GRAE embedding showing the differentiation from astrocytes progenitors to astrocytes. **C** openness of SOX9 TFBSs. **D-E** SOX9, DNAH7 and EFEMP1 expression.

For the PBMC data set, we also analysed the ATAC and GEX modalities. Focusing on the plasmacytoid dendritic cells (pDCs), which are cells responsible for presenting antigens to other immune cells, SEAGALL uniquely identified the RUNX1 and RUNX2 binding motifs (SuppFig6G,H), which are known TFs determining the pDCs lineage^42^. In the same cell type but from GEX data, the TF SOX4 is ranked in the most important features but not in the most differential ones (SuppFig7E), and it is known to be involved in pDCs ontogeny^43^. In addition, we exclusively found CR1 (SuppFig7F) in the explanation of memory B cells, which is known to be necessary for the correct development of this cell type^44^. In the natural killers (NK) and T MAIT cells we uniquely retrieved, respectively, LAIR2 and CD8 (SuppFig7G,H), which are their cell type markers^45,46^.

Combining the features discovered by SEAGALL with motif analysis and manual inspection, we show how SEAGALL can identify several relevant TFBS and genes which are known to be determinants of cell types and their lineages. These TF motifs and genes were not discovered by the classical differential analysis pipeline, showing that our method is able to extract meaningful biological insights which can contribute to the discovery of new determinants of cell identity. In particular, we identified several features, which were not identified using differential analysis, which related to the development and differentiation of cells, suggesting the ability of SEAGALL to capture features which are important in a dynamical state rather than only differences between populations.

## Discussion

In this study we present SEAGALL (Single-cell ExplAinable Geometry-Aware Graph Attention Learning pipLine), a deep learning method based on manifold learning and explainable AI to analyse different modalities of single-cell data. SEAGALL combines a graph-regularised autoencoder (GRAE) and a graph attention network (GAT) together with an explainable artificial intelligence (XAI) method to classify the cells into cell types or states and extract the most important input features for the label prediction (Fig.1). We applied SEAGALL to ten single-cell data sets from three different omics (sc-RNAseq, sc-ATACseq, sc-ChIPseq) and have shown that SEAGALL can consistently understand and explain the cell identity from a different perspective than the classical differential analysis. The combination of a manifold learning method and an autoencoder (GRAE) to reduce the dimensionality of the data has been for the first time extensively applied to the single-cell field. This part of the workflow has been able to reconstruct corrupted data with the highest success (Fig.2), and we also showed that the biological information about cell type was robustly preserved while building the cell-to-cell graph (Fig.3). Therefore, this strategy has revealed an effective and reliable method to build a cell-to-cell graph, which is the final representation of the input data. Moreover, using the geometry aware graph as input to a classifier which applies an attention mechanism to increase the flexibility of the model, the classification performances reach a maximum (Fig.4). The main innovation of SEAGALL is the use of explainable AI to explore the cell type phenotype, making our method highly translational. Often, deep learning has focused on prediction performances, keeping the black box closed and preventing the direct gain of new biological knowledge. Here, we exploit a novel method of graph neural network explainer (GNNExplainer) to open the black box and extract specific, stable and novel features (Fig.5, 6) which drive the cell type classification predictions. Thanks to its user-friendly code and tutorial, we have made our method suitable and useful for real world applications, since it can be directly applied to any count matrix from single-cell data. The deep learning method to learn the data sets ensures that the nonlinearity of the manifold, determined by the complicated gene regulatory networks, is taken into account, while standard approaches based on pca and DA do not. We have applied SEAGALL to several single-cell datasets and have shown that we are able to retrieve TFBSs which are driving factors of cell identity, but that would not have been identified using standard differential analysis pipeline (Fig.6). Finally, SEAGALL can be applied to different single-cell data modalities, such as scATAC-seq, scRNA-seq and scChIP-seq data, reflecting the omic-free hypothesis framework we proposed.

## Methods

### Single-cell RNAseq data processing

Single-cell RNAseq quantifies the abundance of RNA, mainly mRNA, molecules within a cell. For each single-cell the sequencer reads the transcripts that belong to it; hence the output is a raw set of reads which need to be aligned and quantified. For the two human multi-ome data sets (PBMC and brain), raw reads were processed using Cell Ranger Arc 2.0.2 aligning the reads onto the complete human genome (T2T) ^47^. The GEX count matrix of the HSPC data set has been downloaded from^48^. The GEX count matrix of the mouse brain data set has been taken from 10X website^48^. We computed the probability distribution across cells of the number of non zero genes and the number of mitochondrial reads; we filtered out all the cells having a value of one of these variables lying outside the 5% or 95% quantile of their distribution. Similarly, genes present in less than 5% or more than 95% quantile of the cells were moved. Data were library-size normalised. We kept the top 10% highly variable genes. Last, data were log transformed. Differential expressed genes (DEG) between cell types were calculated using the Wilcoxon test. We kept for our analysis the fifty most differentially expressed genes.

### Single-cell ATACseq data processing

Single-cell ATAC-seq is a popular technique to profile chromatin openness at the single-cell level. Typically, When analysing scATAC-seq data, typically the measurements are summarized in a count matrix using the positions of signal enrichment on the genome, called peaks^49^. To construct a count matrix, peaks are called on the pseudo-bulk signal and for each cell and each peak the number of reads that fall into the peak are counted. The structure of the matrix is identical to scRNA-seq, but in the latter case the features are the transcripts.

The reads of the kidney data set^26^ have been processed using Cell Ranger ATAC 2.1.0^2^ and the ones of the PBMC and human brain data sets have been aligned with Cell Ranger Arc 2.0.2; in both cases the reference genome is the T2T human genome^47^. Count matrices were built using episcanpy^12^, given the fragments file and peak file obtained with MACS2^50^. We computed the distribution of the number of features per cell and we filtered out cells having a number of features lower than the 5% quantile or higher than the 95% quantile of this distribution. Cells with lower than 2 for the transcription start site (TSS) enrichment score, and higher than 2 for the nucleosome signal, have been filtered out. Features (peaks) present in lower than 5% or more than the 95% quantile of the cells have been removed. Data were library-size normalised. Only peaks with a variance higher than the one defining the 80% quantile of the variance distribution were kept, with a maximum of 30000 features. Last, data were log transformed.

For the mouse data set, we downloaded the fragments file from 10X database. The fragments file of the HSPC data set was downloaded from the original publication^48^. Before building the count matrix and filtering, following the procedure described above, we called peaks using MACS2^50^.

The choice to use quantile as thresholds is motivated by the fact that this an automatic and fully reproducible method to apply quality controls on the data: instead of inspecting matrix by matrix and apply to every case a different threshold without being able to fully motivate the choice, the quantile computation takes into account the specific properties of the data set, ensuring the same rigidity on the filtering.

sc-ChIPseq experiment count matrices were downloaded from the original publication^27^. In this case the features are windows, that are constant size (50kb) intervals spanning the whole genome. We processed those data as peaks since the processing does not rely on any peaks-specific assumption. Differential open peaks or windows between cell types are calculated with the Wilcoxon test. We kept for our analysis the fifty most differentially open peaks or windows.

### Cell type annotation

For the HSPC^48^, kidney^26^ and breast cancer^27^ data the cell type annotation is provided from the authors. The cell type annotation of the mouse brain is taken from^48^ and it is based on marker genes. Human PBMC has been manually annotated following the muon tutorial^51^. Mouse brain has been manually annotated with marker genes and the procedure is shown in our github. Each data set consists of a different number of cell types (SuppTable1,2).

### Embedding and graph construction

Once the count matrices are cleaned we use GRAE to build the cell-to-cell graph. First, PHATE is applied as a manifold learning method; it is able to capture both global and local structure of the data and embed it into a smaller representation with arbitrary dimension. The loss function of the autoencoder, which is the mean squared error (MSE) between original and reconstructed space, is then regularised by adding a term which increases if the AEs latent space differs more from the PHATE embedding. In other words, the total loss function *L* is composed by two terms: a reconstruction term *L*_*r*_ and and a regularisation term *L*_*g*_

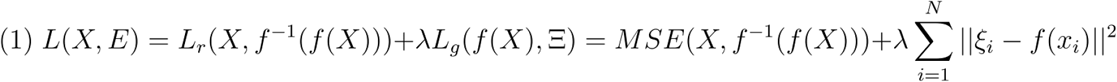

where is *X* a set of *N* data points such that *x*_*i*_ ∈ ℝ^d^, Ξ is the PHATE embedding of *X* such that ξ_*i*_ ∈ ℝ^p^ with *p* < < *d, f* and *f* ^−1^ are, respectively, the encoding and decoding function. The dimension of the latent representation varies for each count matrix to fit the data set complexity and it is set as the cubic root of the number of features.

Within the latent space, pairwise euclidean distance between cells is computed and then a k-NN graph is built, with *k* = 15. k-NN graph had already been used in the literature as input graph for GNNs^52^ but there is also a technical motivation that led us to a constant degree network: building a correlation or distance based graph is intrinsically problematic; after computing pairwise distances or correlations a cut-off is applied to the maximum distance or minimum correlation. Each node may have any number of neighbours in the interval [0, *N* − 1]. We tested this possibility and it turns out the resulting graph is extremely dense (SuppFig8), which may lead to nonsensical connections and makes the training of the GNN extremely time and energy demanding.

The graph is the final representation of the data set, which contains the connectivity pattern and the geometry of the input manifold.

### Cell type classification with GNN

Graph neural networks are a particular type of neural network able to process data with a graph structure. GNNs take as input a graph *G* = (*V, E*), where *V* ∈ ℕ is the set of nodes and *E* ⊆ *V* × *V* is a set of edges, also known as links, between nodes. Each node can have a feature vector that defines the properties of the nodes. In our context, the feature vector is the gene expression or the chromatin openness vector. From a point cloud perspective, the embedding of each point is a function of the point itself and the points close to it. Let *G* = (*V,E*) be an undirected graph containing *N* vertices, *x*_*i*_ ∈ ℝ^*d*^ is the initial representation of node *i*, 𝒩_*i*_ = {*j*∈ *V* | (*j, i*) ∈ *E*} the neighbours of node *i*, then the first layer of the GNN will create a new representation of the node 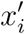 according to (2), known as the message passing equation^53^

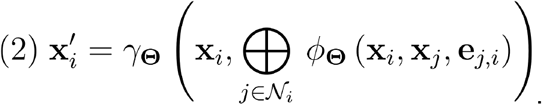

GNN is a broad class of neural networks which rely on equation (2). We decided to apply a more refined version of the base message-passing layer called “GAT”^20^ which applies an attention mechanism to the embedding function, meaning that the model learns the importance of each feature and link, following

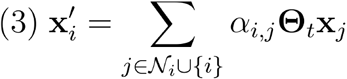

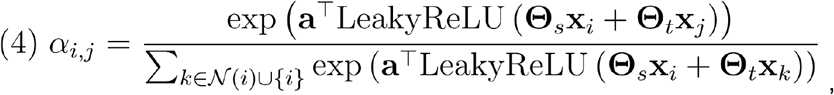

where 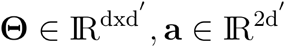 are learned parameters, ⊕ is any differentiable and permutation invariant function such as sum or mean, and *γ*_**Θ**_ and *ϕ*_**Θ**_ are differentiable functions such as MLPs.

Thus, the key property of GNNs is the ability to create latent representations of a local neighbourhood rather than a single point. The rationale behind the choice of GNN relies on this property: we want to have a local analysis of each cell aiming for a local ensemble study, rather than treat them totally independently. Each count matrix with its own graph is given as input to the GNN classificator; the target output is the cell type of each node, which is defined as explained in the “cell type annotation” paragraph.

Our specific model consists of a graph neural network with two layers, first one to create a latent representation of the input and second one which performs the classification task. The dimension of each layer is defined with hyperparameter optimization (HPO)^54^ case by case. The model is trained with Adam^55^ optimizer with learning rate and weight decay estimated with HPO.

### GAT explanation

Once the model is trained an XAI method is applied to it. We choose to apply “GNNExplainer”^22^, which is a model agnostic method. It creates a graph and a feature mask to spot which is the minimum set of features and edges of each node sufficient to predict the class. We assume that the nature of our data defines a real function *f* that labels objects, nodes in the case of GNN, representing cells in our context. The GNN model Φ receives as input a graph *G* and a feature vector *X* as explained in the previous paragraph. In practice, Φ learns a probability *P*_Φ_(*Y*|*G,X*) with *Y* random variable for the classes {*C*_*i*_} *i* = 1,..,*C* representing the probability of nodes to belong to each of the classes. After the training, the model is fixed and it will be used to make predictions. The crucial point of the explainer is the fact that each node has a computation graph *G* and certain node features *X* that completely determine all the information that are necessary to predict 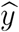 at certain node *v*. Given a node *v*_*i*_ the explainer finds the subgraph *G*_*s*_ ⊆ *G* and the associated features *X*_*s*_ = {*x*_*i*_ | *v*_*j*_ ∈ *G*_*s*_} that maximise the probability of having seen the prediction 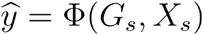 where Φ is the trained GNN. Indicating as *MI* the mutual information function and *H* the entropy function, the GNNExplainer solves the following problem

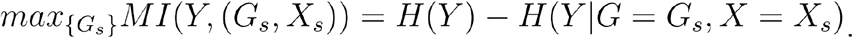

MI quantifies the variation in the prediction probability when the graph and the features are *G*_*s*_ and *X*_*s*_ instead of *G* and *X*, with the feature vector constrained to be much smaller than the original one. In practice, for each node we obtain the features ranked by their importance. Since we are interested in the cell types explanations, we average the feature importance of all the nodes belonging to the same class to obtain the most relevant features for each label.

### Topological and variational autoencoder models

Whereas GRAE^19^, PeakVI^11^ and scVI^10^ are released as packages, we had to implement the models for topological and variational autoencoder. Both the methods are based on the same architecture, which consists of one input layer, one hidden layer, with dimension equal to the square root of the input length, and a latent space, with dimension equal to the cubic root of the input layers. The varying size of the layers are important to account for the data set complexity. The best values of dropout, learning rate, weight decay, weight of topological regularisation and signature of the p-norm (for TopoAE) and weight of Kullback-Leibler divergence for VAE, have been estimated using HPO implemented with the *optuna* package^56^. Each HPO consists of 25 runs to explore the parameters within defined intervals (SuppTable8). We used a subset of count matrices to explore the HPO and we then applied the same parameters for each matrix.

### Input data reconstruction

To test the robustness of the AEs we measured their ability to reconstruct the input data after corruption. To corrupt the data, we applied an increasing dropout from 10% to 50% of the features in linear steps of 10% to each count matrix and we trained each model with the corrupted data. All the models have been trained with the same patience (20) and maximum number of epochs (300). We used 85% of the data for training and 15% for validation. When applying dropout the choice of the removed features is random, therefore it may happen that we remove some features particularly important for a specific model but not for another. To make sure our results are not biased by this factor, we repeat this experiment ten times varying the features to drop out at each level. For each run, for each level of dropout and for each model we measured the MSE between the original data (the not corrupted one) and the model reconstructed data. We then computed the average MSE for each level and model and the uncertainty of the mean.

### Graph homogeneity

After applying each dimensionality reduction method as described in the previous paragraph (GRAE, TopoAE, VAE, PeakVI, scVI and pca), but without dropout, to each one of the count matrices, we computed the k-NN graph (k=15) from their latent spaces. For each cell we computed how many different cell types are found in its neighbourhood. We divided this value by both the number of neighbours (15) and the number of cell types (different for each data set) to create a score called heterogeneity. Last we computed the homogeneity as 1-heterogeneity.

### Classification and XAI experiments

To test the quality of each embedding method we used their latent space to build the cell-to-cell k-NN graphs (k=15) and we gave the graphs as input to a graph neural network node classifier. We tested the combination of the six dimensionality reduction methods (GRAE, TopoAE, VAE, PeakVI, scVI and pca) and two different GNN: GAT and GCN (see *Cell type classification with GNN* paragraph for the details of the models). We run each combination fifty times. Each training started with a different random seed to make sure that the models do not always start from the same point in the parameter space. Both GAT and GCN are trained for 250 epochs with a patience of 20 epochs. The data sets have been splitted into train, validation and test sets with a ratio of, respectively, 70%, 10% and 20%. Before training the classifier we run a 25 steps HPO study to select the best values (SuppFig9) of each hyperparameter of the GNN, within defined ranges (SuppTable9). After each training we applied the explainer for 200 epochs and saved the fifty most important features for each label, i.e. for each cell type. Accuracy, precision, recall and F1 score have been computed in the standard way using *sklearn*^57^; the specificity of the explainer is defined as one minus the average intersection of the top fifty most relevant features across cell types. Stability is defined as the average intersection of the explanation for the same cell type across different runs of the classifier and the explainer.

## Data and code availability

Code and tutorial for SEAGALL are available at https://github.com/gmalagol10/seagall.git

Data and notebook to reproduce all the results are available at https://github.com/gmalagol10/seagall/tree/main/reproducibility.

## Acknowledgments

We thank Samuele Firmani for the insightful discussion about the evaluation of the models. We thank Vera Manelli for the important help in the interpretation of motif analysis. We thank Federica Tosato for the suggestions about visualisation and graphics. We thank Gaia Fontana for drawing the logo of the tool.

G.M. is supported by the Helmholtz International Lab Causal Cell Dynamics (InterLabs-0029) - Grant support from the Initiative and Networking Fund of the Hermann von Helmholtz-Association Deutscher Forschungszentren e.V.-. G.M is also supported by the Helmholtz Association under the joint research school “Munich School for Data Science — MUDS and by German Research Foundation project ID 213249687–SFB 1064. We thank the BMC Bioinformatics Core Facility for providing access to their HPC cluster. Figures have been made with the help of Biorender ^58^.

## Contributions

G.M., M.C.T. and G.W. designed the study and conceived the algorithm. G.M. implemented the algorithm. P.H. provided code and helped the implementation of it. A.D. annotated the human brain data set. G.M. and M.C.T. wrote the manuscript with help from G.W. and additional inputs from all co-authors. All authors reviewed and approved the manuscript.

## Supplementary Figures

**SuppFig1:**
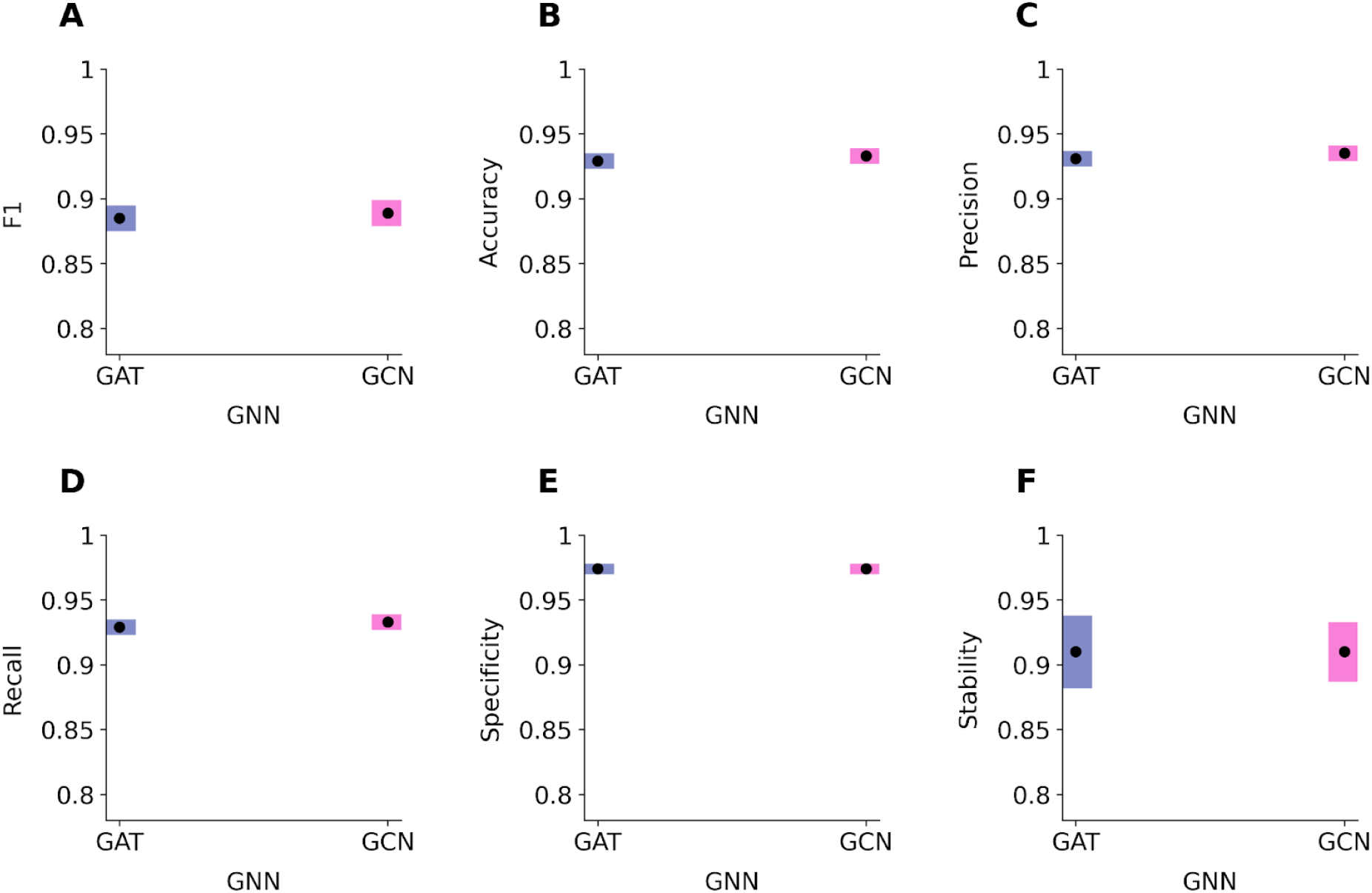
**A-D** classification of the two GNN architectures. Black dots indicate the mean over all the runs and the different embedding methods and the bar represent three times the uncertainty on the mean. **E-F** specificity (left) and stability (right) of the two GNN architectures. In none of the six metrics here represented we can appreciate a significant difference between the two models.

**SuppFig2:**
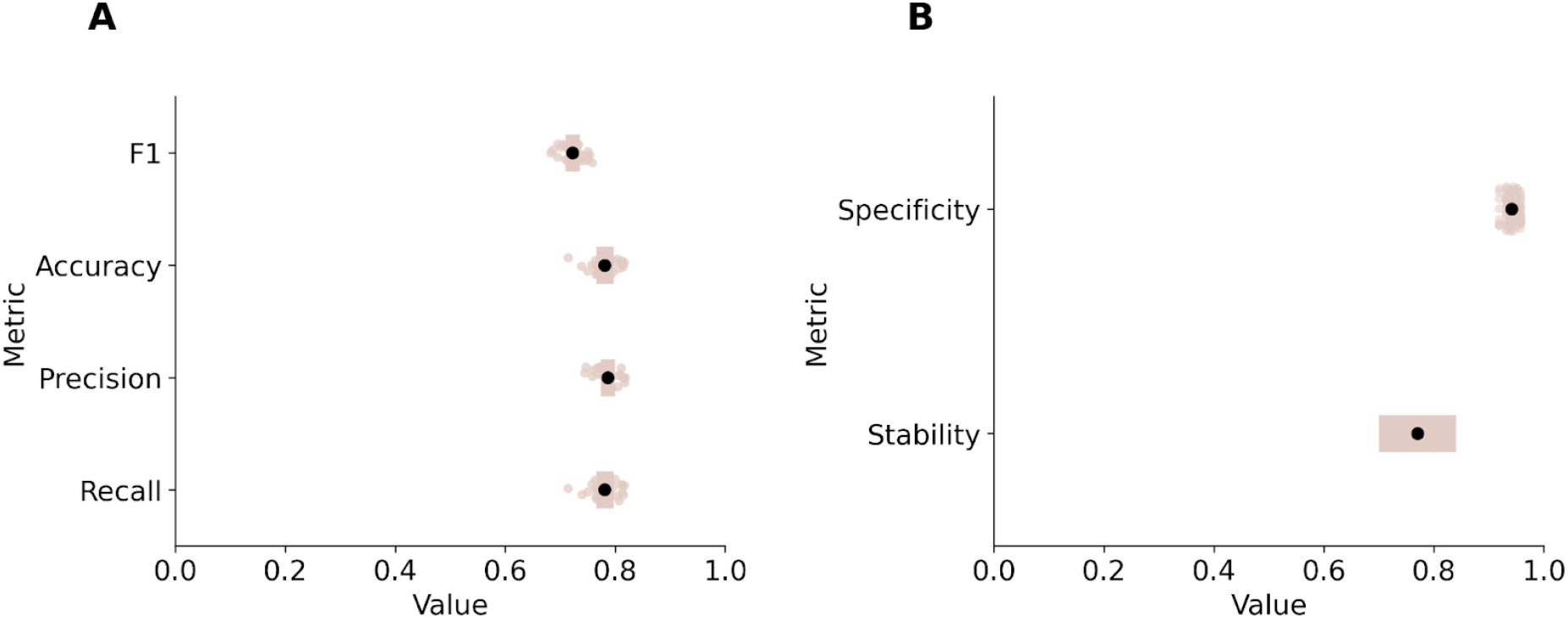
**A** classification performances of the final model on the scChIP-seq. **B** stability and specificity of the explainer on the scChIP-seq data set.

**SuppFig3:**
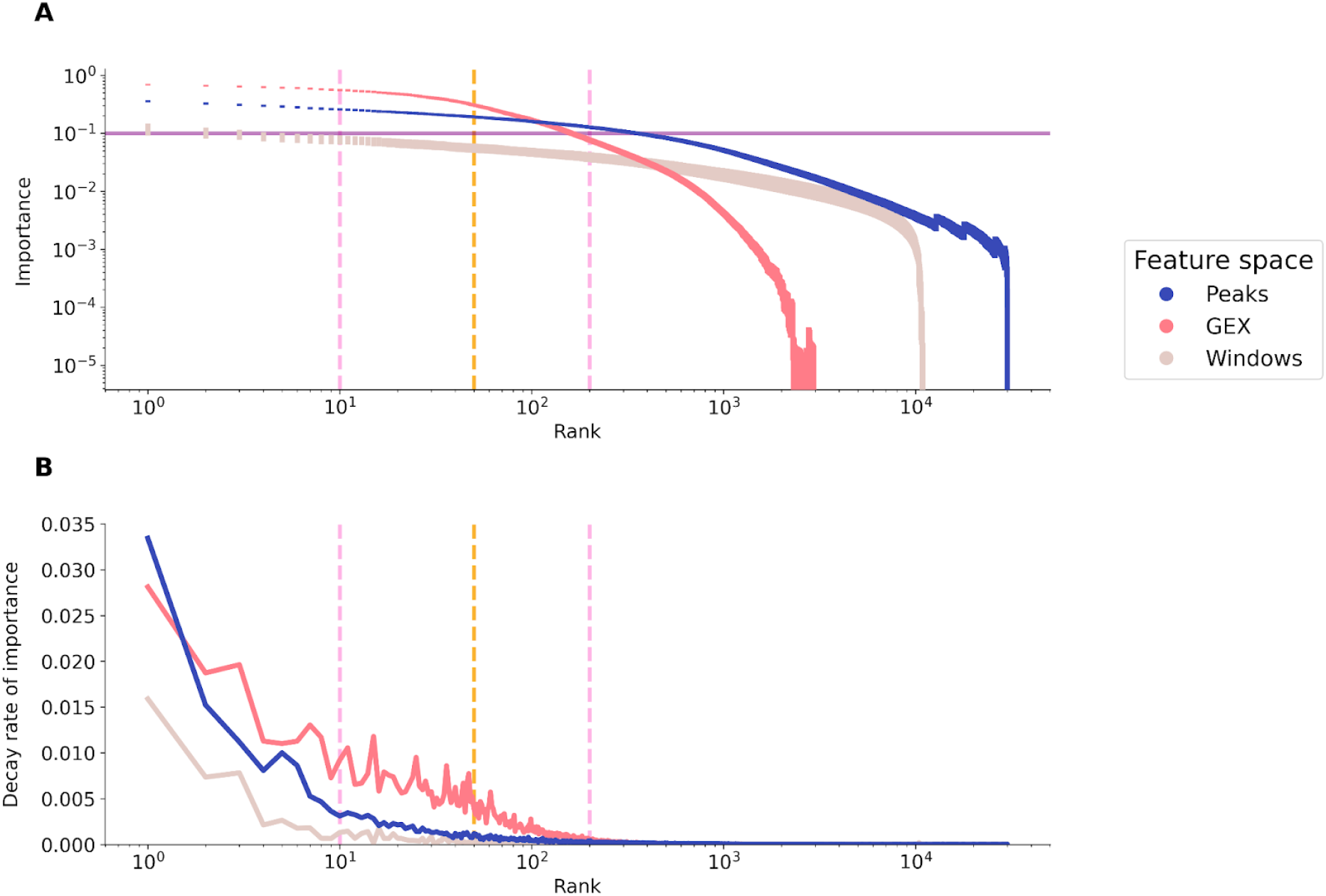
**A** full range rank-importance distribution of the features for each feature space. **B** full range derivative of importance distribution of the features for each feature space.

**SuppFig4:**
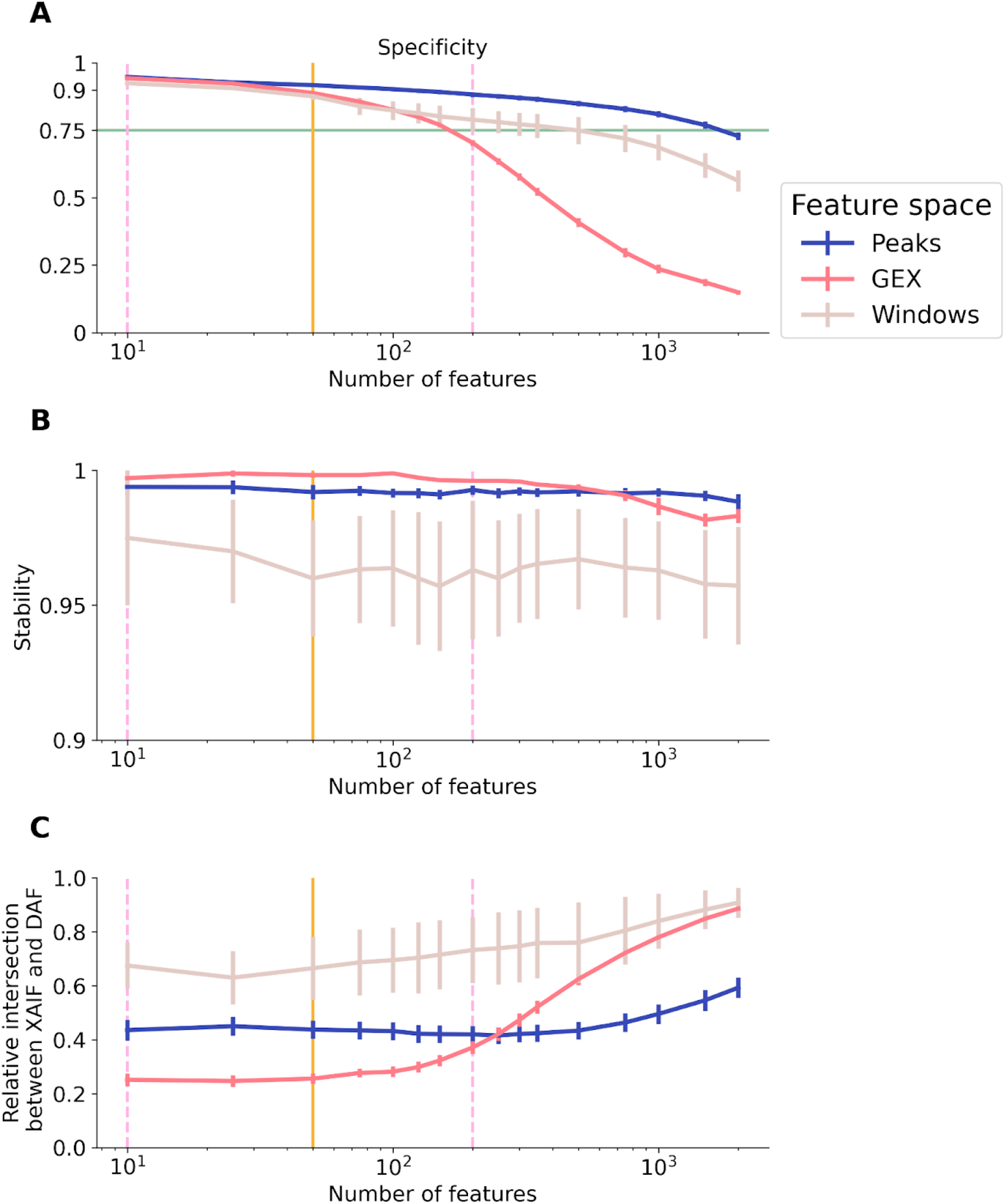
**A** specificity of the explanations varying the number of features we asked the model to keep. Each color represents one feature space; the mean value at each step is the average of the specificity for each cell type in each data set for each after running the explainer fifty times. The error bar height represents the standard deviation of the mean. **B** stability of the explanation computed as described in A. **C** Similarity of the XAIF to the DAF measured in terms of relative overlapping.

**SuppFig5:**
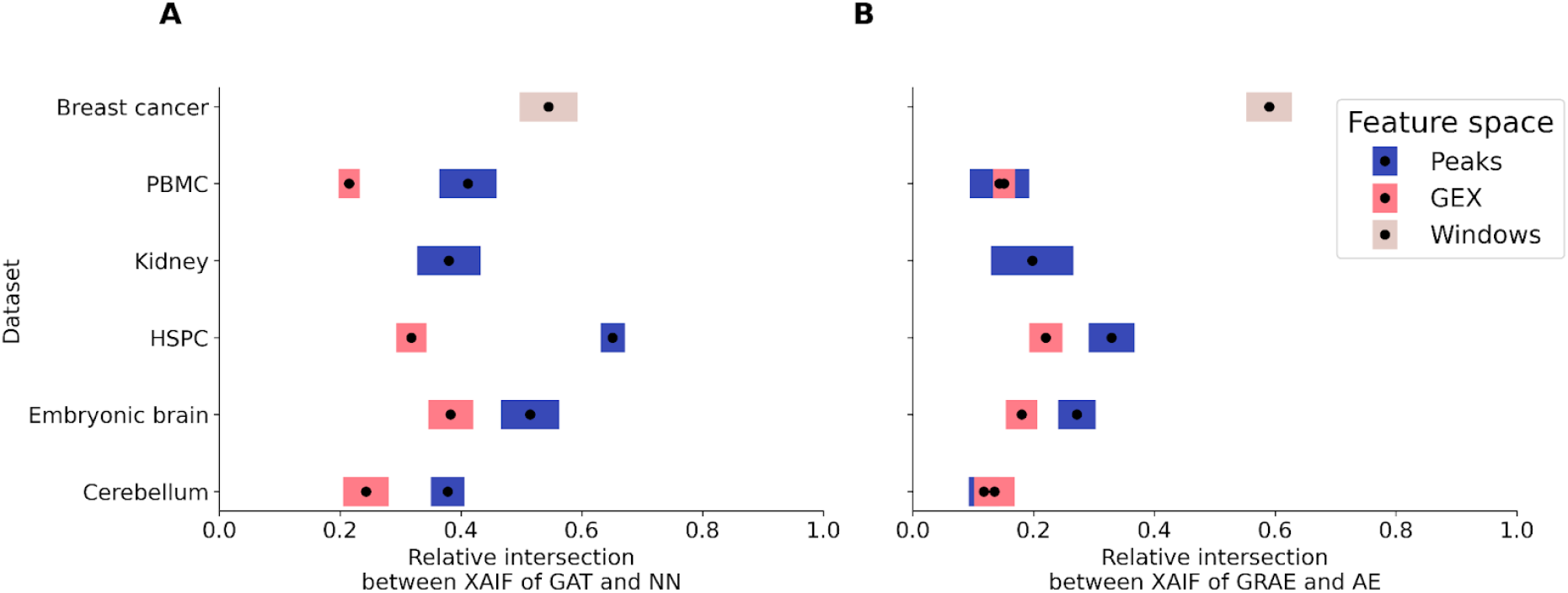
**A** Difference between the XAIFs of GAT and a normal NN without any graph information. **B** Difference between the XAIFs of GRAE versus AE for graph construction, applying the same GAT. Removing either geometry or attention leads to different explanations.

**SuppFig6.**
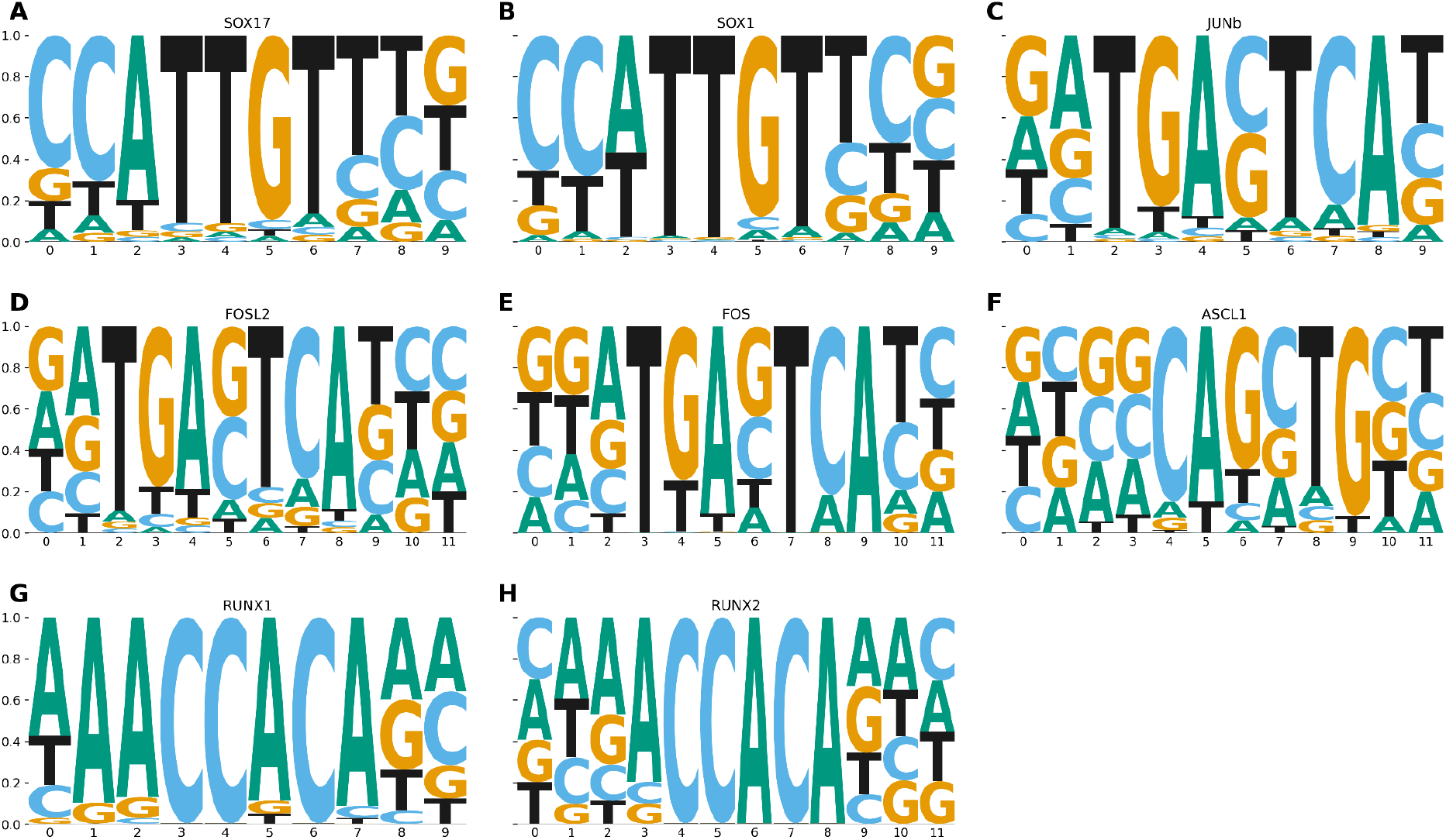
**A-B** motif visualisation

**SuppFig7:**
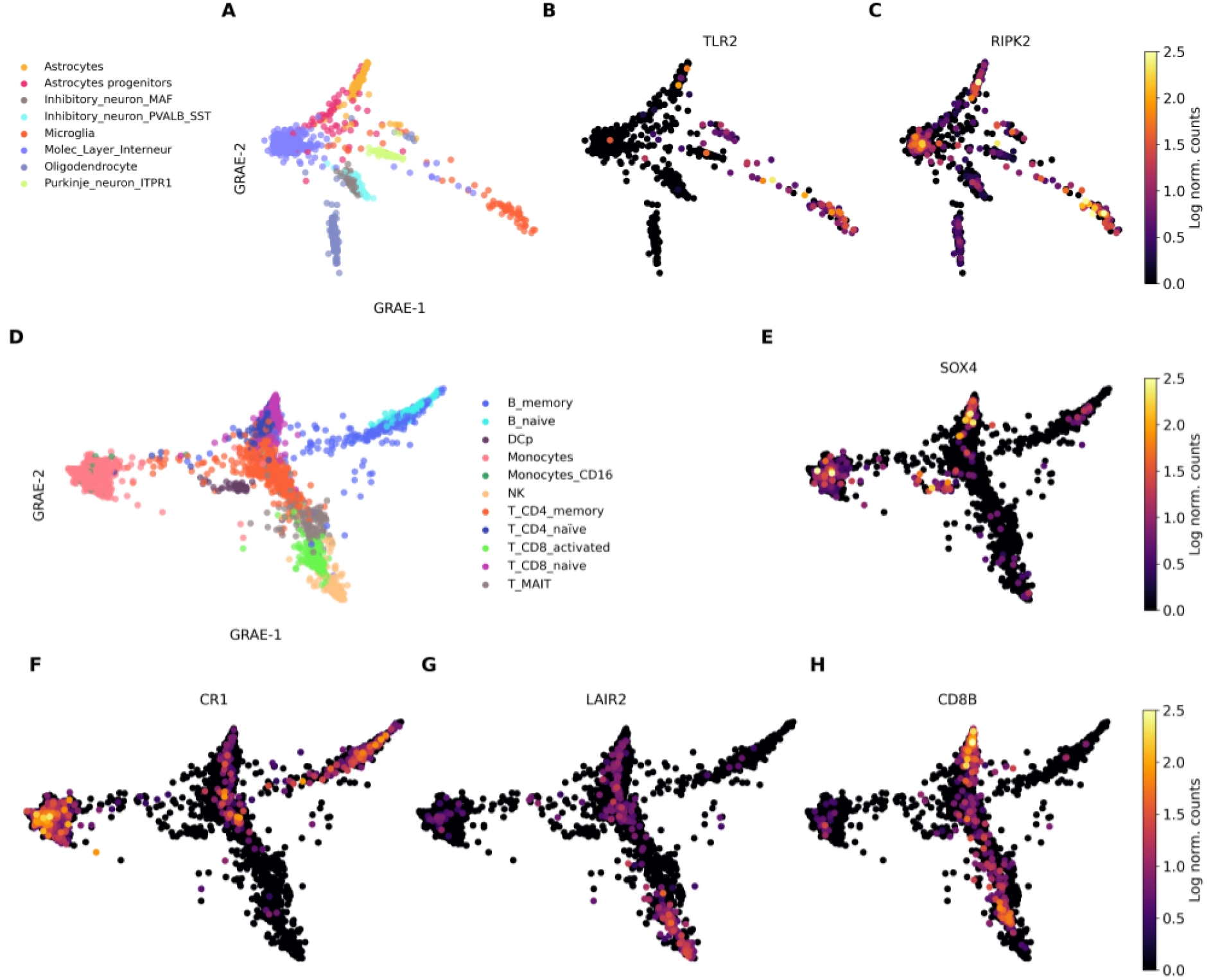
**A** GRAE embedding of the sc-ATAC of the human brain data set. **B-C** expression of TLR2 and RIPK2 in normalised log counts. **D** GRAE embedding of the GEX of the PBMC data set. **E-H** expression of SOX4, CR1, LAIR2 and CD8 (sub unit B) in normalised log counts.

**SuppFig8.**
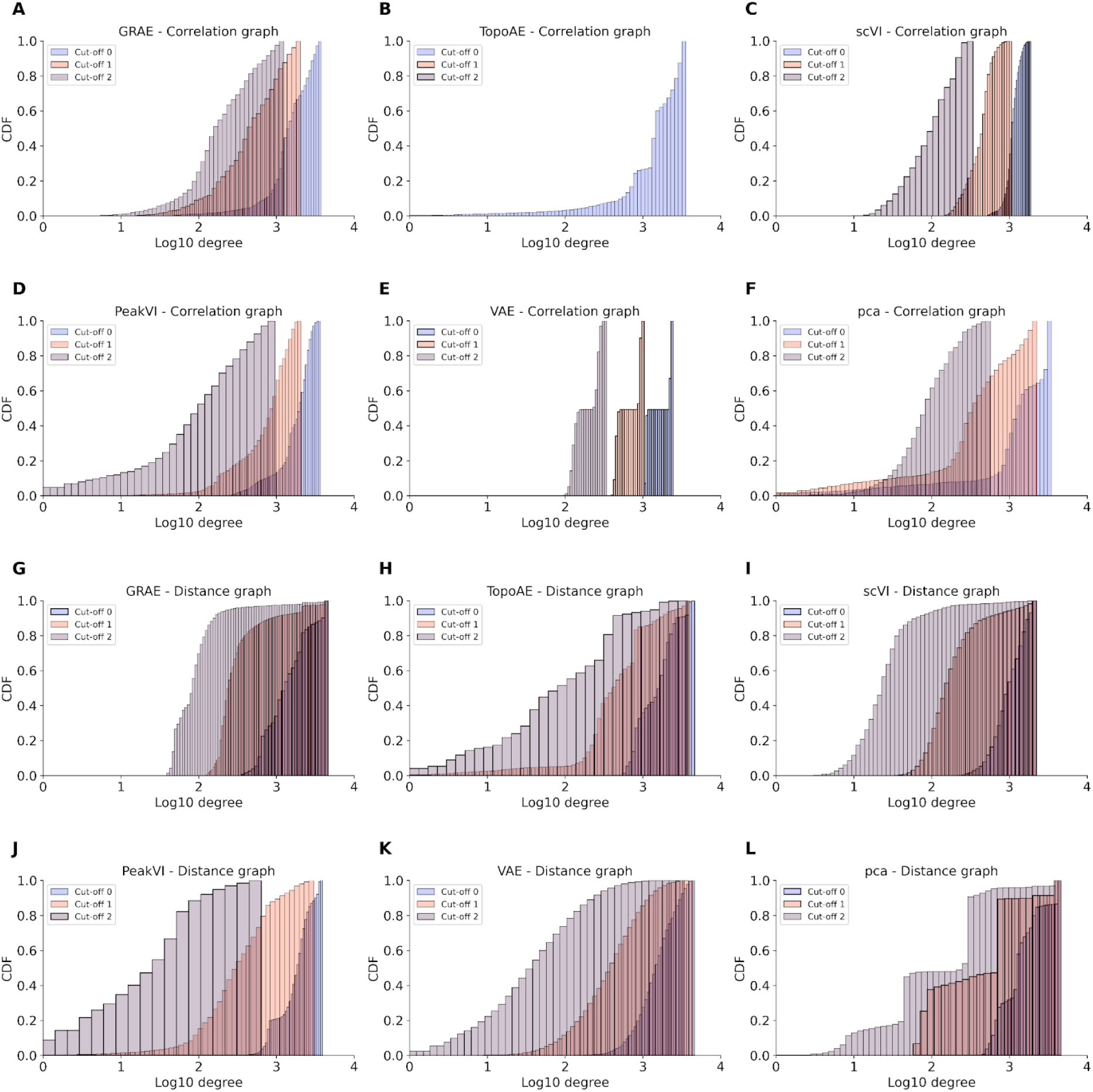
**A-F**: cumulative distribution function of nodes degree of the correlation graph varying the latent space (across panel) and applying different cut-off on the minimum correlation between nodes to keep the link. **G-L** show the same information but based on euclidean distance computed in the latent space of the embedding method. In all the cases, the graphs are very dense making the edges meaningless and the computation very time and energy demanding.

**SuppFig9:**
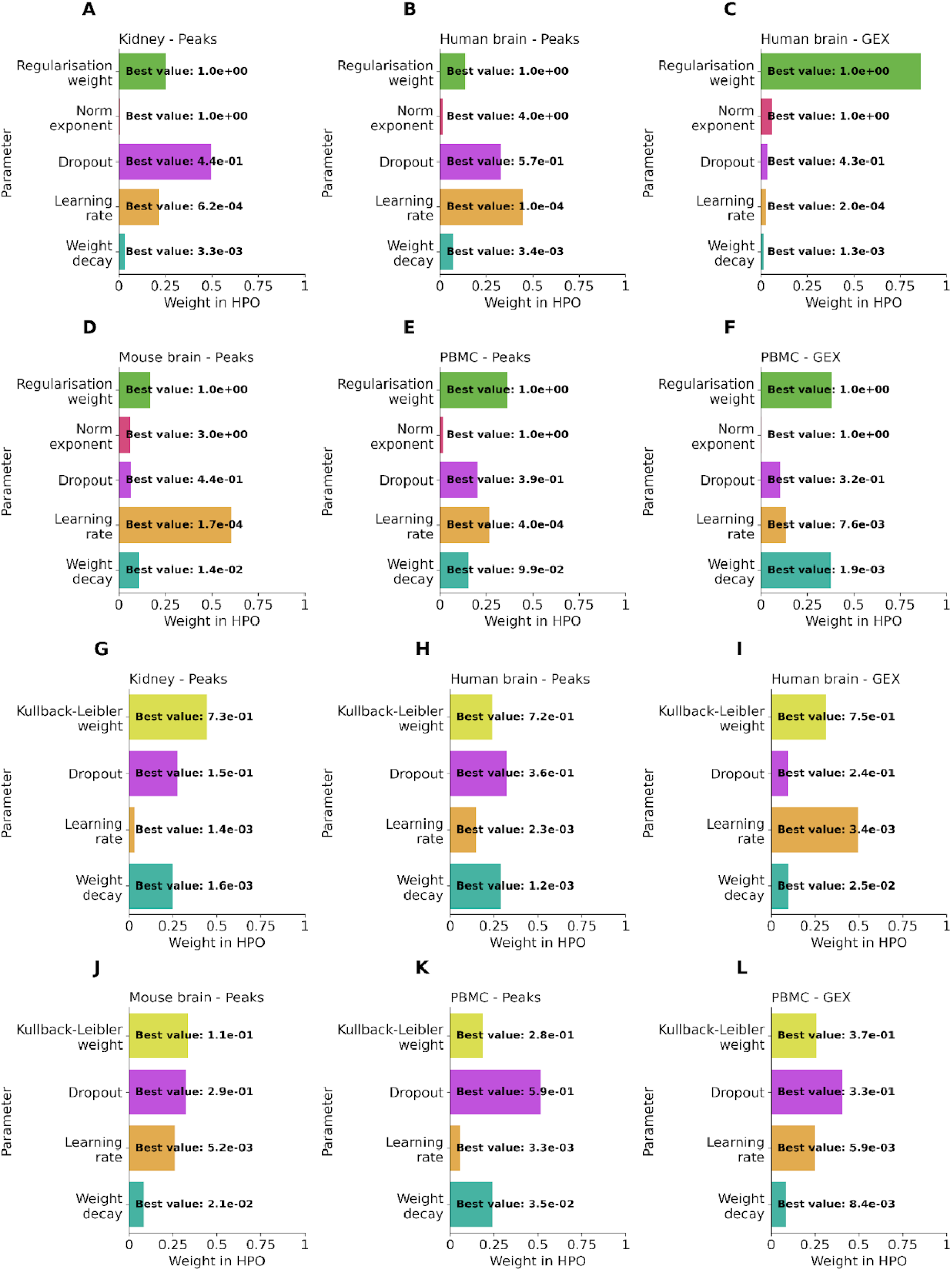
impact (weight) of each hyperparameter in the optimization of either TopoAE or VAE. The bold annotations indicate the best values of the hyperparameter, i.e. the ones used in all the experiments involving TopoAE (**A-F)** and VAE (**G-L**). Each panel corresponds to one data set. The final value of each hyperparameter is the average across the six experiments.

## Supplementary Tables

**Supplementary table 1:** cell type composition of each data set.

|  | Human brain | Breast cancer | PBMC | HSPC | Mouse brain | Kidney |
| --- | --- | --- | --- | --- | --- | --- |
| 0 | Molec_Layer_Interneur | HBCx-95 | Monocytes_CD16 | HSC | Subplate | Endo |
| 1 | Microglia | HBCx-95-Ca paR | T_CD8_naive | LMPP | Deeper_Layer | PT |
| 2 | Oligodendrocyte | HBCx-22 | B_naive | MPP | Upper_Layer | CDPC |
| 3 | Purkinje_neuron_layer | HBCx-22-Ta mR | B_memory | MEP | IPC | Immune |
| 4 | Inhibitory_neuron_PVALB_SST |  | NK | Prog_B | V_SVZ | LOH |
| 5 | Purkinje_neuron_ITPR1 |  | T_MAIT | Prog_DC | RG_Astro_OPC | CDIC |
| 6 | Inhibitory_neuron |  | T_CD4_naive | Erythrocyte | Ependymal_cells | Podo |
| 7 | Purkinje_neuron_FOXP2 |  | DCm | GMP |  | DCT |
| 8 | Astrocyte |  | Monocytes | Granulocyte |  |  |
| 9 | Inhibitory_neuron_MAF |  | T_CD4_memory | Prog_MK |  |  |
| 10 | Astrocyte_progenitor |  | T_CD8_activated | Platelet |  |  |
| 11 |  |  | DCp |  |  |  |

**Supplementary table 2:** number of features and cells of each count matrix.

| Data set | Feature space | Number of features | Number of cells |
| --- | --- | --- | --- |
| HSPC | GEX | 2689 | 10389 |
| HSPC | Peaks | 30000 | 10350 |
| Kidney | Peaks | 25901 | 4624 |
| Human brain | GEX | 3003 | 2441 |
| Human brain | Peaks | 17886 | 2726 |
| PBMC | GEX | 2761 | 5700 |
| PBMC | Peaks | 12781 | 6372 |
| Mouse brain | GEX | 2237 | 2721 |
| Mouse brain | Peaks | 29114 | 3027 |
| Breast cancer | Windows | 10908 | 4636 |

**Supplementary table 3:** average homogeneity of each k-NN graph computed from the latent space of the AEs. The uncertainty is three times the standard deviation of the mean. Bold numbers highlight the method achieving the best performances; multiple bold numbers indicate that the methods performances fall within the uncertainty. The empty spaces represent the forbidden combinations of AE and feature spaces, which are GEX with PeakVI and peaks with scVI.

|  | Kidney<br>Peaks | Human brain<br>Peaks | Human brain<br>GEX | Mouse brain<br>Peaks | PBMC<br>Peaks | PBMC<br>GEX |
| --- | --- | --- | --- | --- | --- | --- |
| GRAE | **0.9905±0.0003** | **0.9911±0.0004** | **0.9917±0.0003** | **0.9849±0.0004** | **0.9909±0.0002** | **0.9921±0.0002** |
| TopoAE | 0.9875±0.0004 | 0.9894±0.0005 | **0.9915±0.0004** | 0.9799±0.0006 | 0.9896±0.0002 | 0.9907±0.0002 |
| VAE | 0.9634±0.0005 | 0.9666±0.0007 | 0.9668±0.0006 | 0.9570±0.0008 | 0.9671±0.0004 | 0.9671±0.0004 |
| PeakVI | 0.9882±0.0003 | **0.9909±0.0004** |  | **0.9846±0.0005** | 0.9903±0.0002 |  |
| scVI |  |  | **0.9921±0.0003** |  |  | **0.9922±0.0002** |
| pca | 0.9766±0.0004 | 0.9771±0.0008 | **0.9921±0.0003** | 0.9701±0.0006 | 0.9856±0.0003 | **0.9923±0.0007** |

**Supplementary table 4:** classification and explanations performances of the two GNN architectures. GCN and GAT achieve the same performances. Values are the mean over all the runs and the using GRAE as embedding method. The uncertainty is three times the standard deviation of the mean.

|  | Accuracy | F1 | Precision | Recall | Specificity |
| --- | --- | --- | --- | --- | --- |
| GAT | 0.929±0.006 | 0.89±0.01 | 0.93±0.01 | 0.93±0.01 | 0.974±0.004 |
| GCN | 0.933±0.006 | 0.89±0.01 | 0.94±0.01 | 0.93±0.01 | 0.974±0.004 |

**Supplementary table 5:** performances of the GAT classifier varying the embedding methods. For each method are reported the mean across 50 runs and three times the uncertainty on the mean. GRAE outperforms all the methods except scVI in terms of accuracy, F1 score, precision and recall. In bold is highlighted the best performing method, the second best one is underlined. See SuppTable4 and SuppTable5 for the performances divided by feature space.

|  | Accuracy | F1 | Precision | Recall | Specificity | Stability |
| --- | --- | --- | --- | --- | --- | --- |
| GRAE | <u>0.929±0.006</u> | <u>0.885±0.010</u> | <u>0.931±0.006</u> | <u>0.929±0.006</u> | <u>0.974±0.004</u> | **0.91±0.03** |
| TopoAE | 0.857±0.013 | 0.778±0.015 | 0.865±0.012 | 0.857±0.013 | <u>0.975±0.004</u> | **0.91±0.03** |
| VAE | 0.369±0.034 | 0.33±0.04 | 0.48±0.04 | 0.369±0.034 | 0.960±0.007 | 0.81±0.04 |
| PeakVI | 0.909±0.007 | 0.842±0.013 | 0.909±0.007 | 0.909±0.007 | **0.981±0.001** | **0.89±0.05** |
| scVI | **0.956±0.006** | **0.928±0.005** | **0.959±0.006** | **0.956±0.006** | **0.975±0.007** | **0.92±0.02** |
| pca | 0.734±0.032 | 0.665±0.033 | 0.781±0.023 | 0.734±0.032 | **0.980±0.001** | 0.84±0.04 |

**Supplementary table 6:**
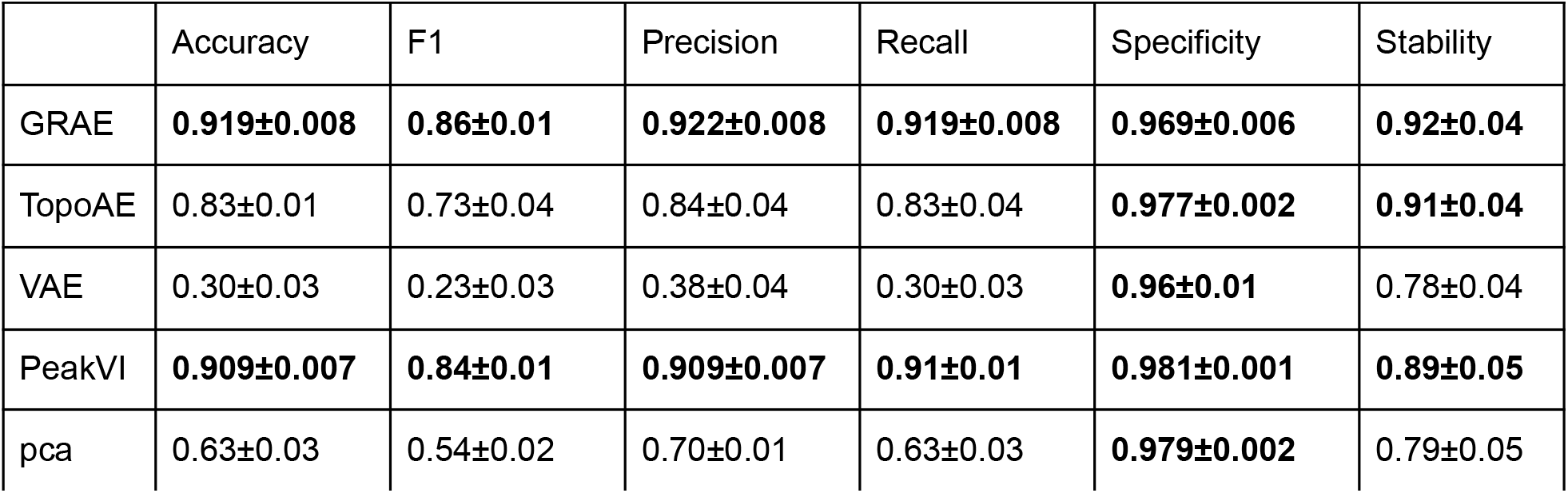
classification and explanation performances of the embedding methods as listed in Table but only for peaks count matrices.

**Supplementary table 7:** classification and explanation performances of the embedding methods as listed in Table but only for GEX count matrices.

|  | Accuracy | F1 | Precision | Recall | Specificity | Stability |
| --- | --- | --- | --- | --- | --- | --- |
| GRAE | **0.95±0.01** | **0.929±0.007** | **0.950±0.009** | **0.95±0.01** | **0.9819±0.0007** | **0.89±0.04** |
| TopoAE | 0.91±0.02 | 0.87±0.02 | 0.92±0.02 | **0.92±0.02** | **0.97±0.01** | **0.90±0.03** |
| VAE | 0.50±0.05 | 0.53±0.03 | 0.67±0.03 | 0.50±0.05 | **0.9668±0.0009** | **0.87±0.04** |
| scVI | **0.956±0.006** | **0.928±0.005** | **0.959±0.006** | **0.956±0.006** | **0.975±0.007** | **0.92±0.02** |
| pca | **0.950±0.007** | **0.921±0.006** | **0.953±0.007** | **0.950±0.007** | **0.9822±0.0006** | **0.92±0.02** |

**Supplementary table 8:** minimum and maximum explored values during HPO of the parameters of AEs.

|  | Regularisation weight | P for the P-norm | Weight decay | Learning rate | KL weight | Dropout |
| --- | --- | --- | --- | --- | --- | --- |
| Min value | 1 | 1 | 0.0001 | 0.0001 | 0.1 | 0.1 |
| Max value | 30 | 5 | 0.1 | 0.1 | 0.9 | 0.7 |
| AE | TopoAE | TopoAE | TopoAE<br>VAE | TopoAE<br>VAE | VAE | TopoAE<br>VAE |

**Supplementary table 9:** minimum and maximum explored values during HPO for the GNNs parameters.

|  | Hidden dimension | Number of heads | Weight decay | Learning rate |
| --- | --- | --- | --- | --- |
| Min value | 32 | 4 | 0.0001 | 0.0001 |
| Max value | 256 | 12 | 0.1 | 0.1 |

## Extended Tables

**Extended Table 1:**
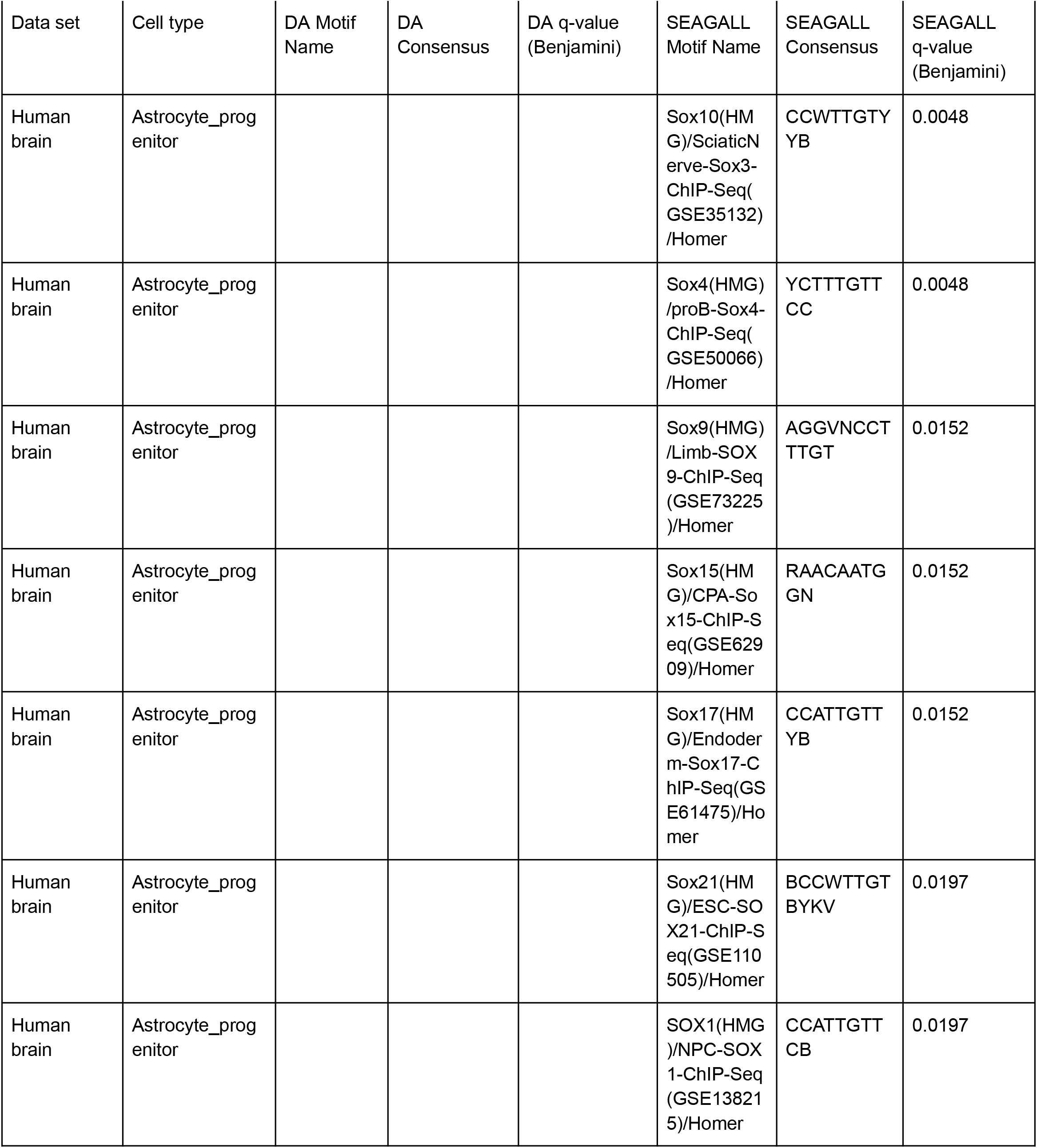
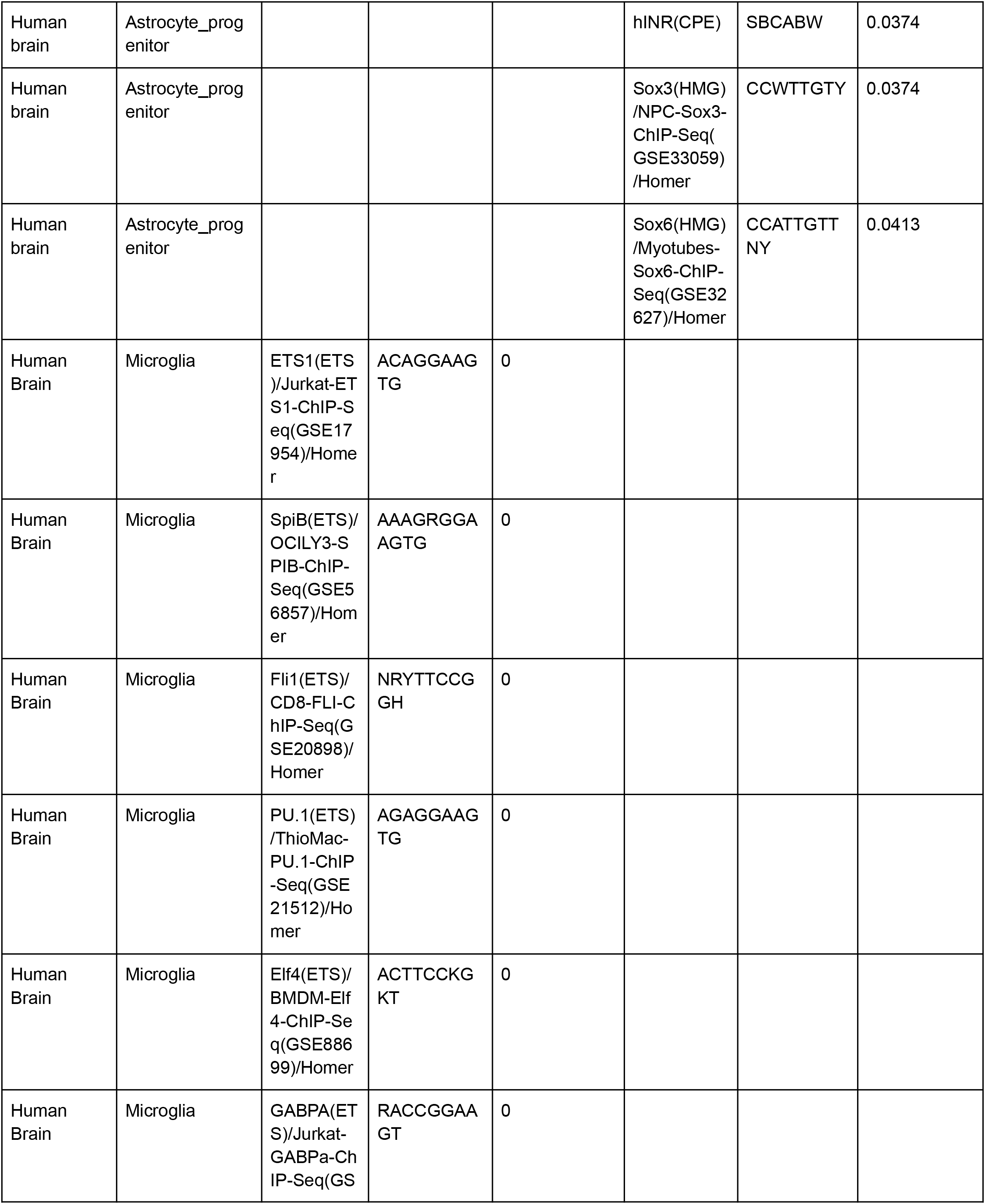
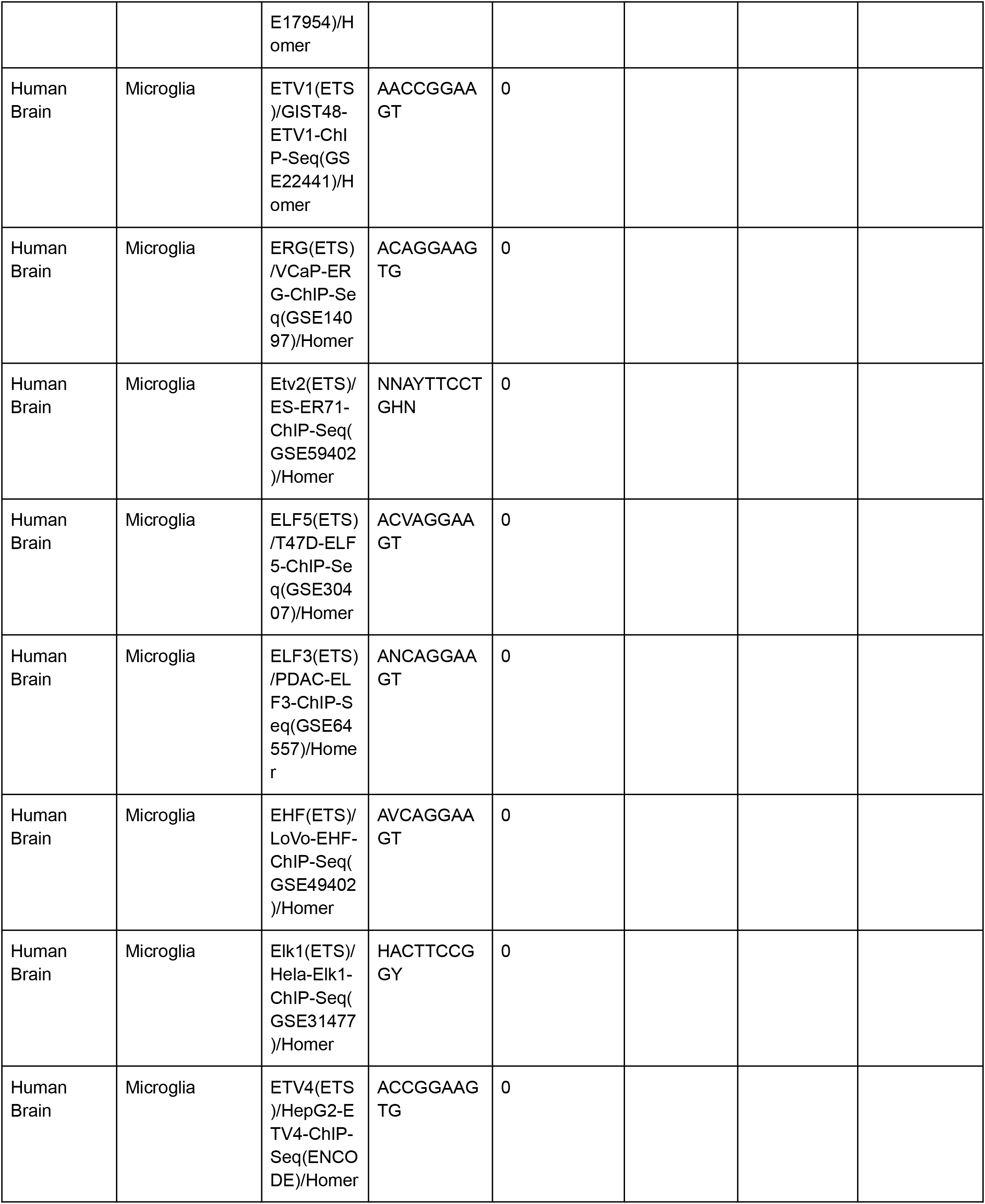
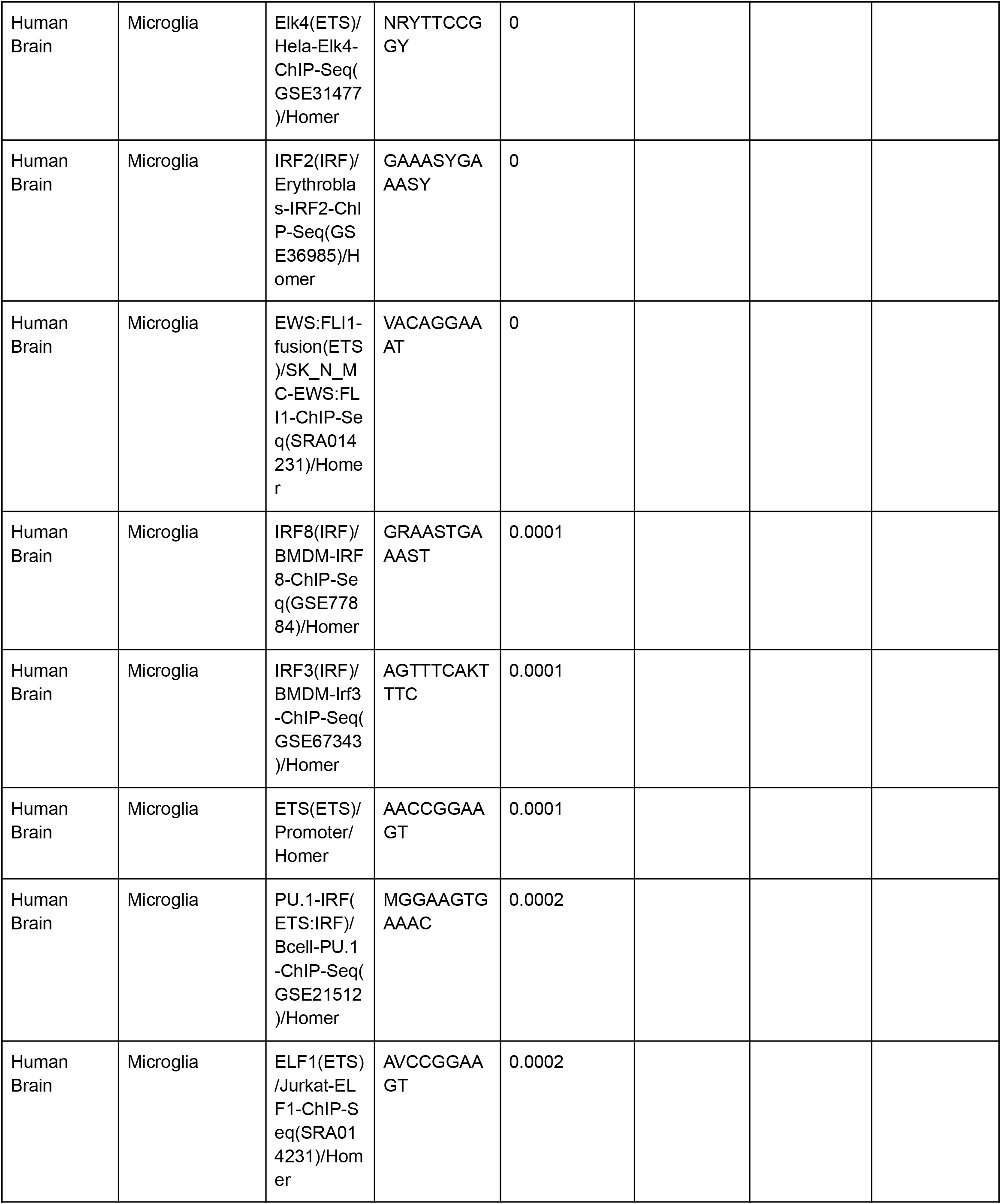
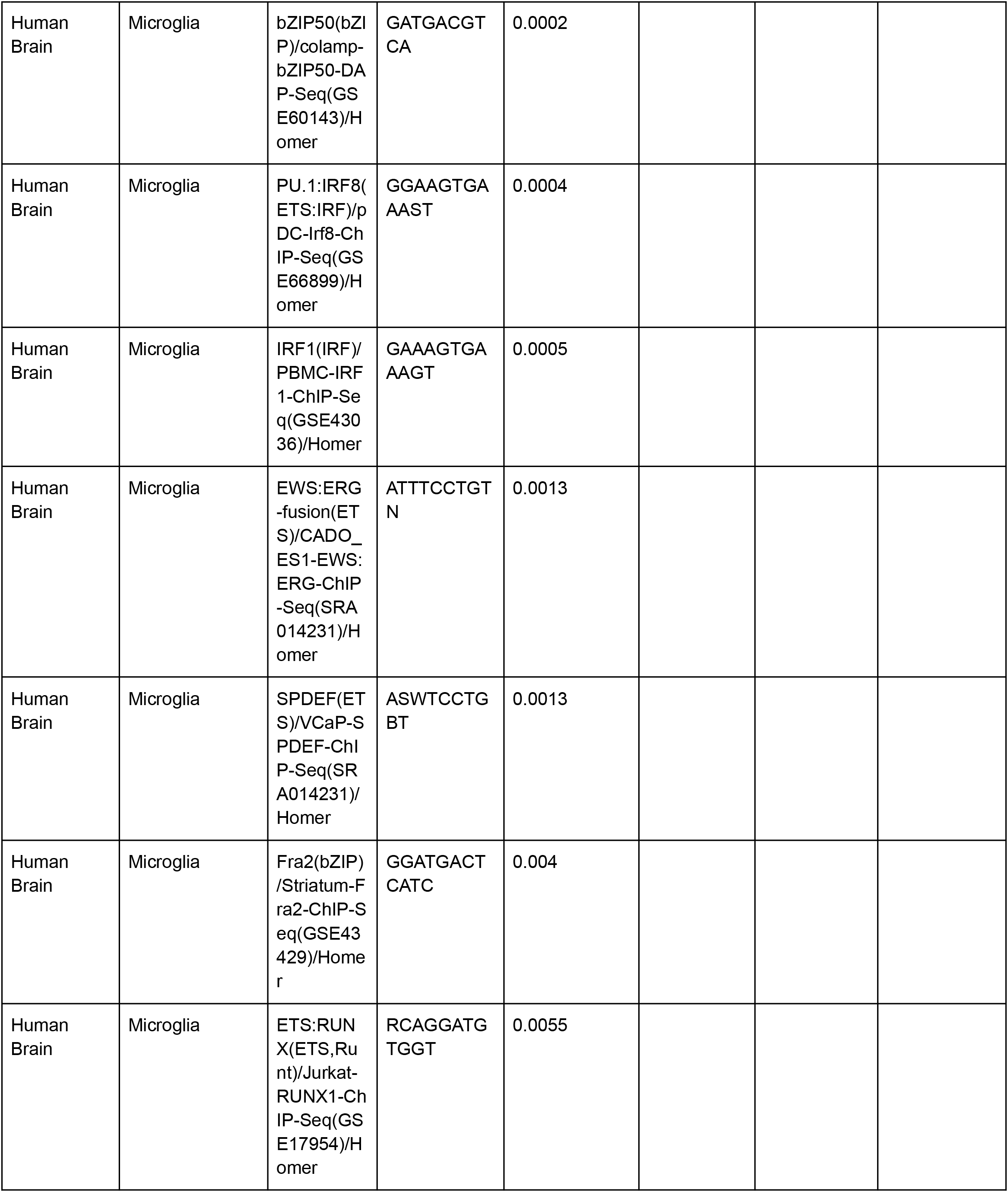
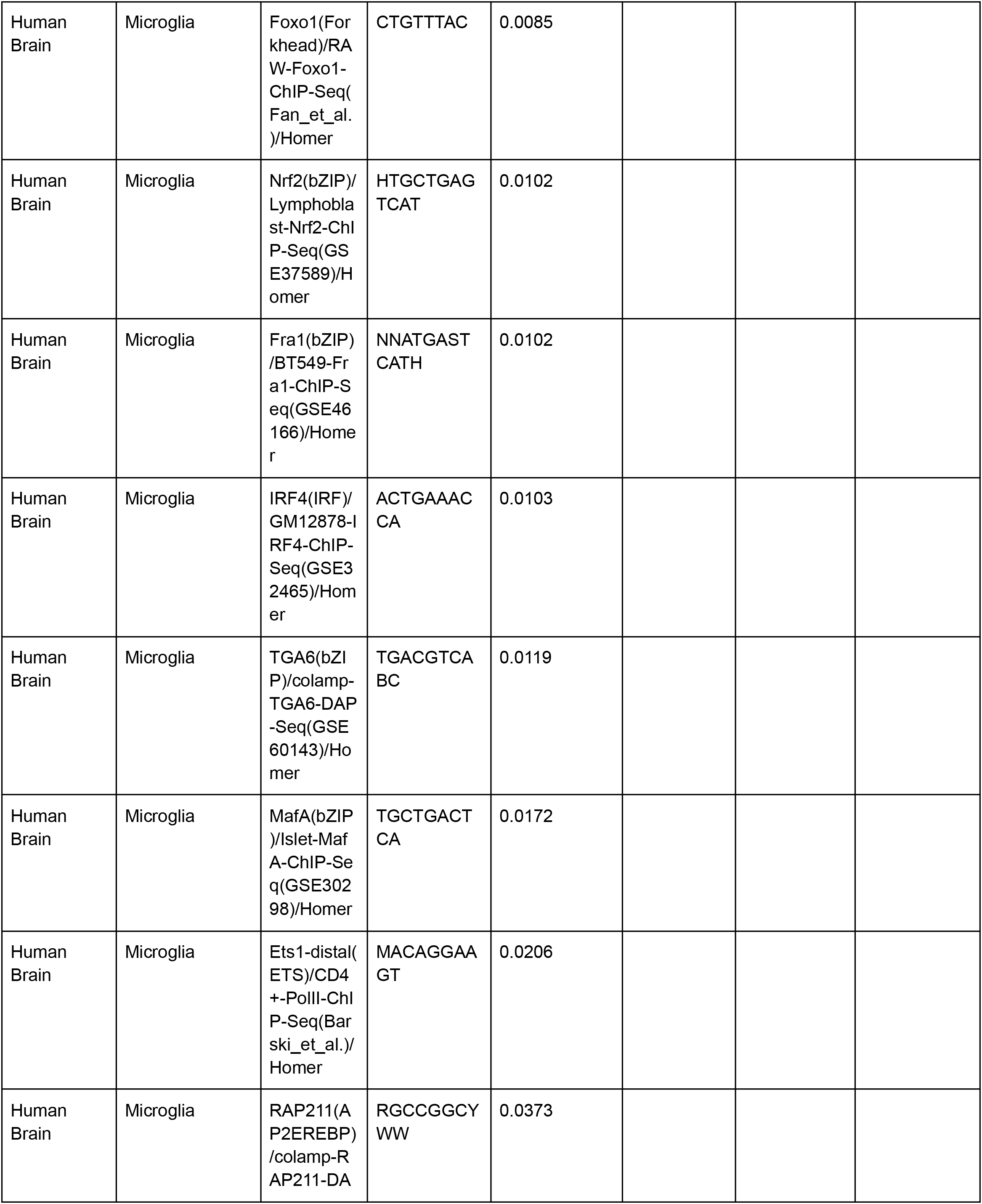
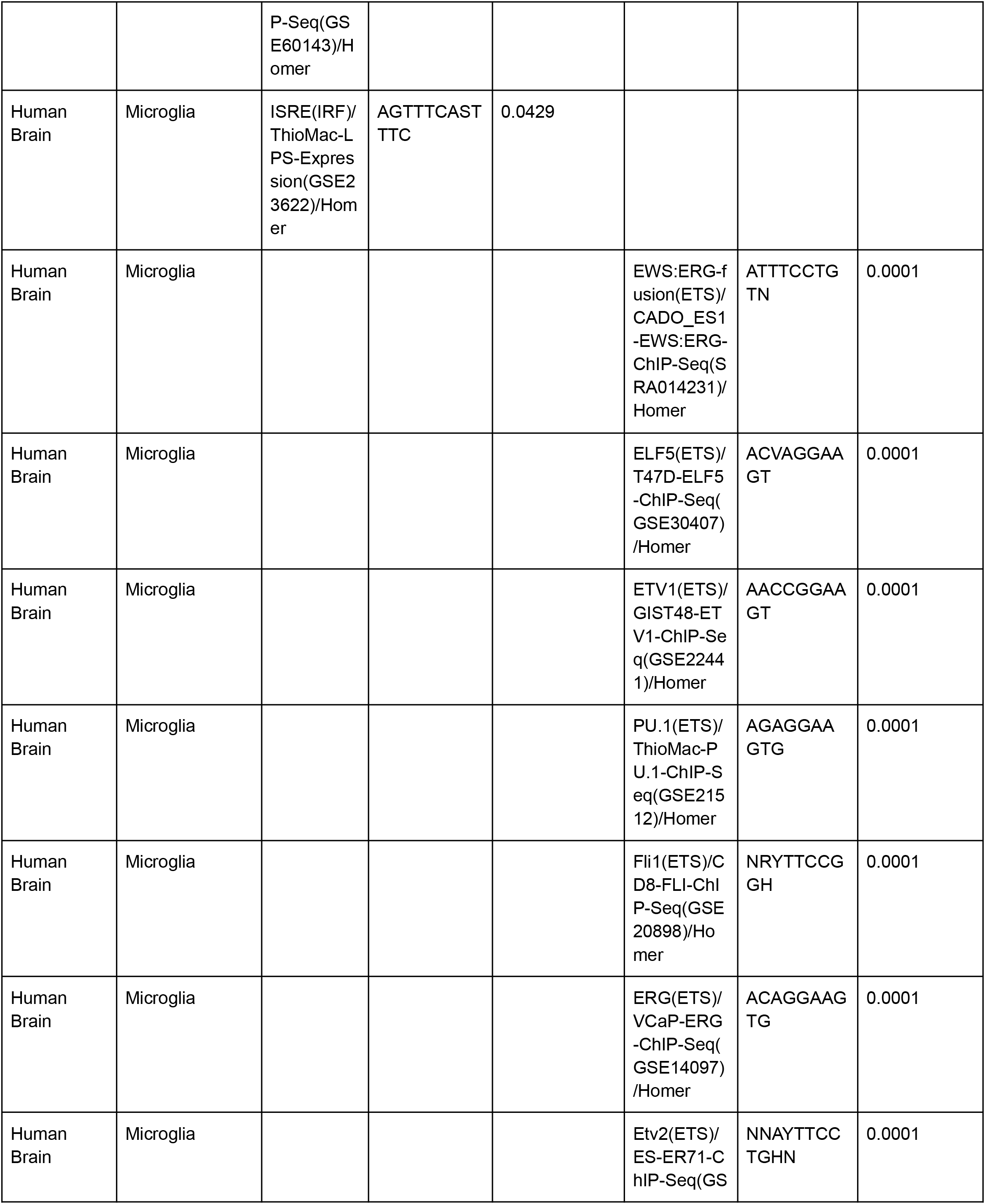
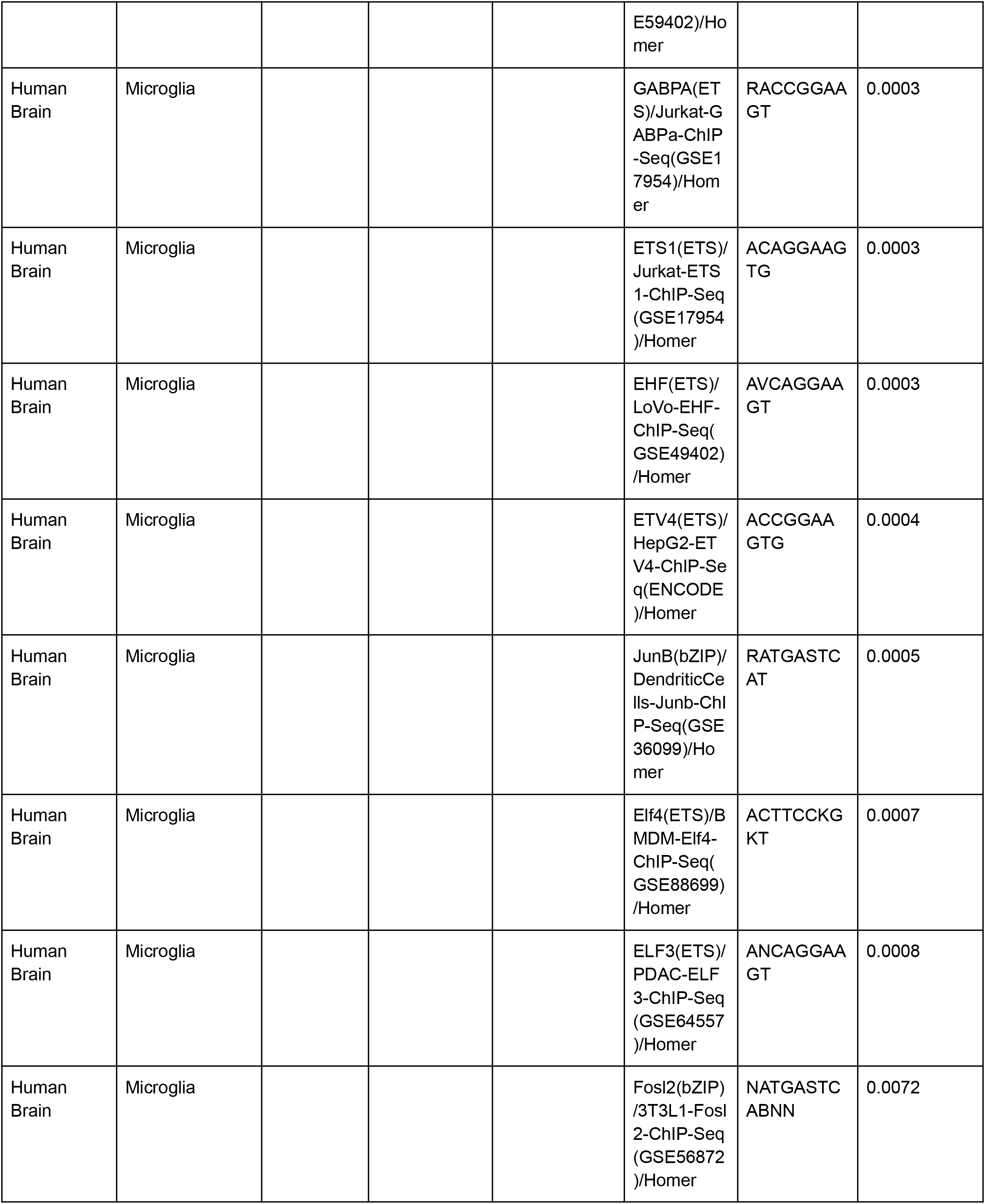
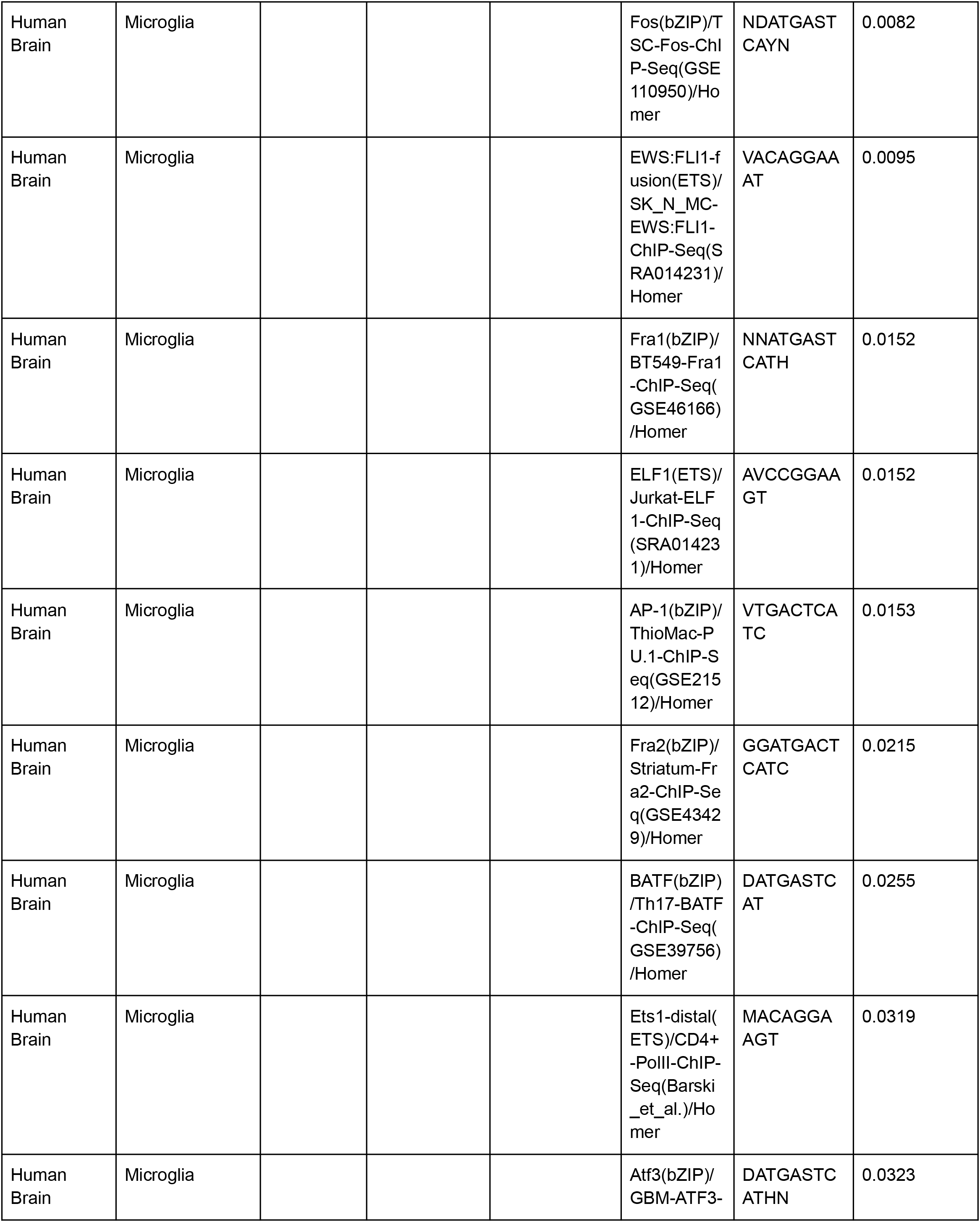
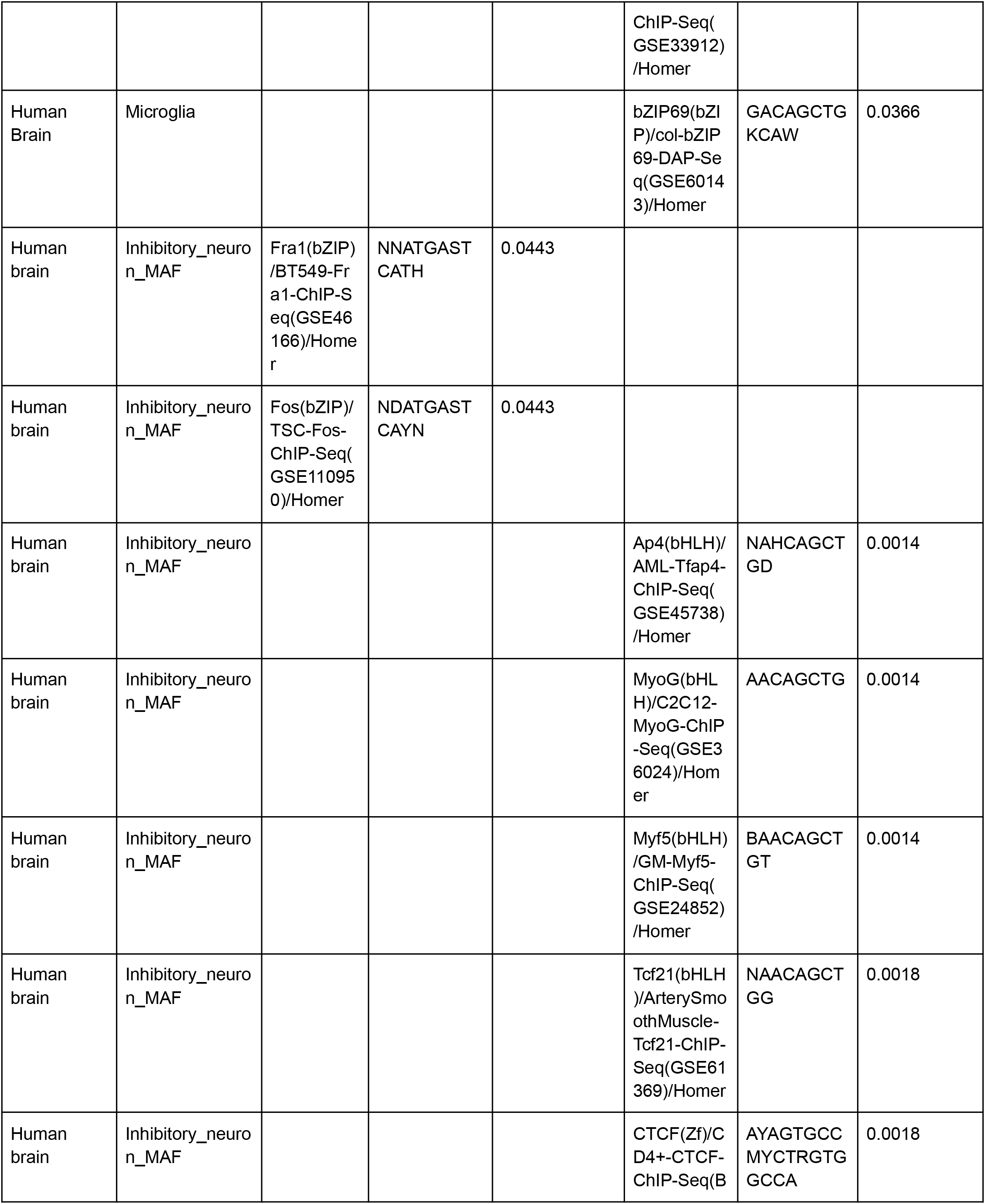
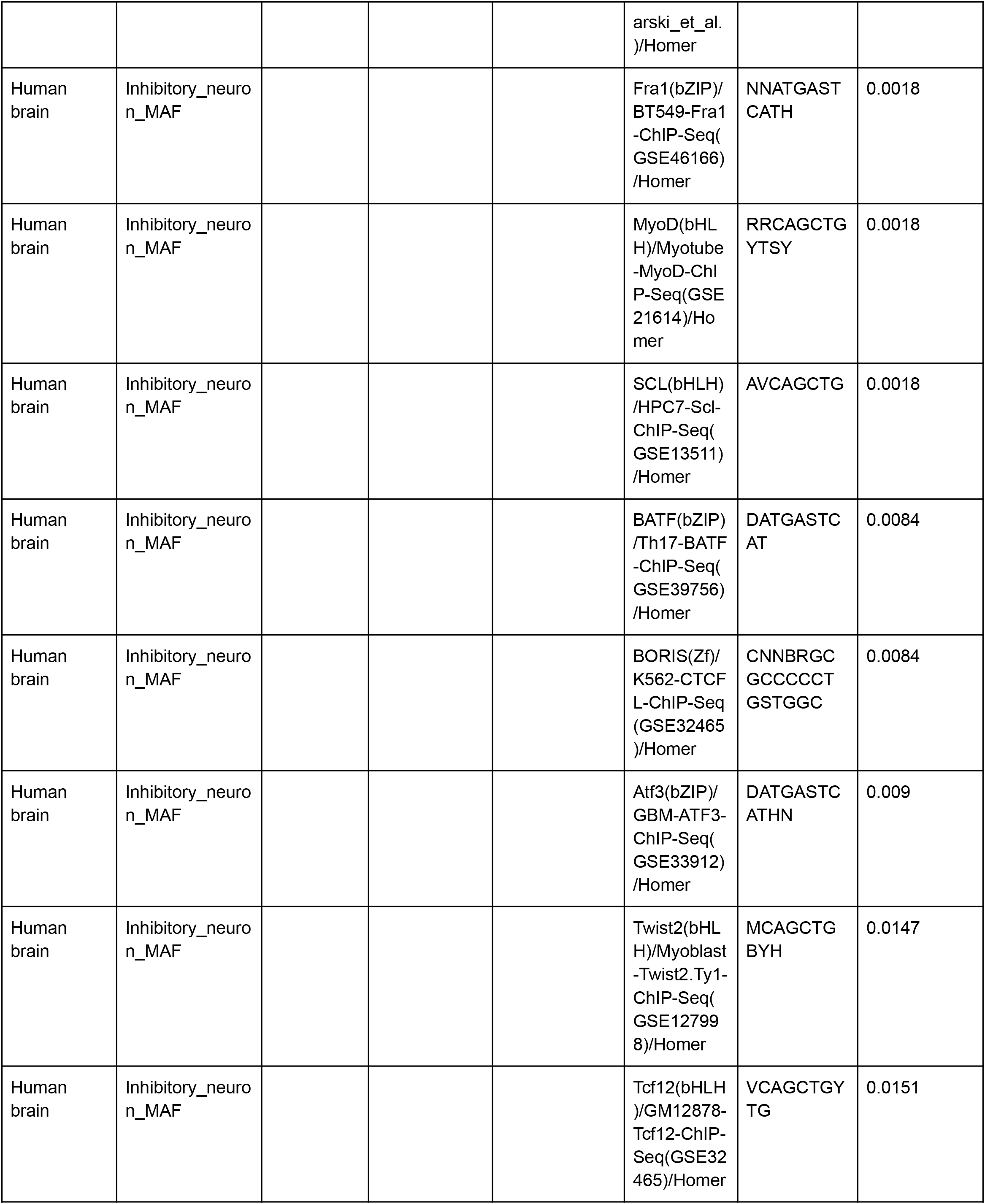
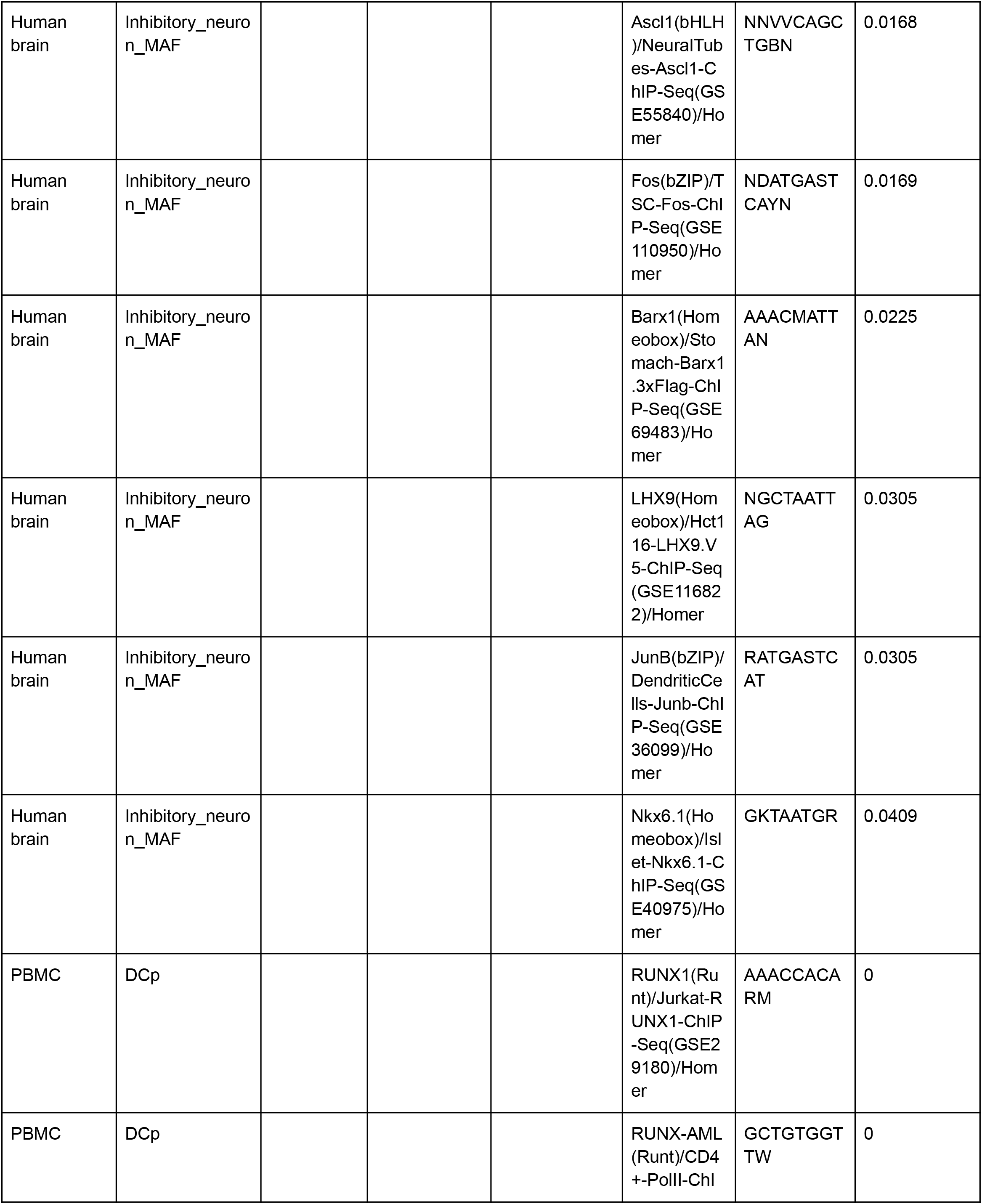
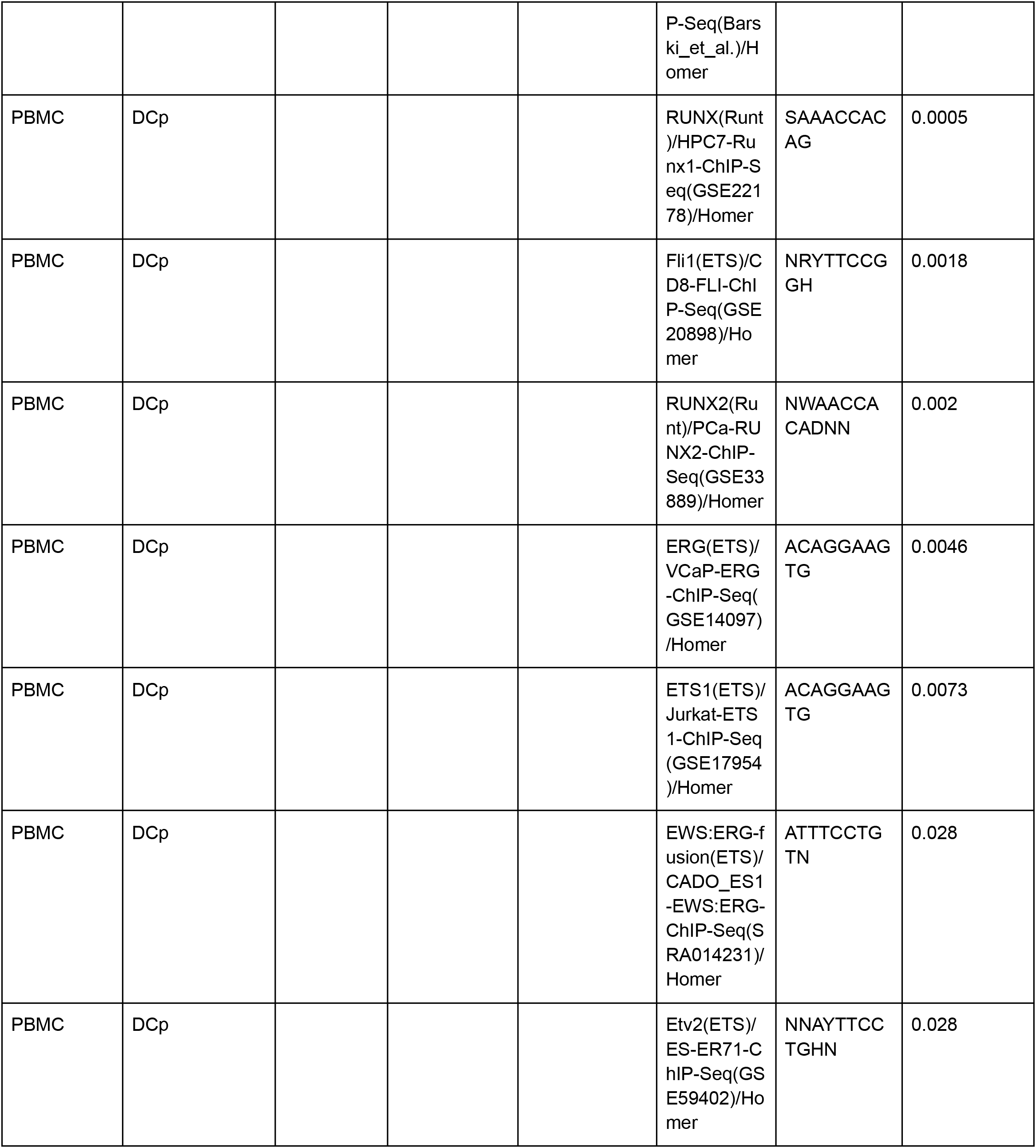
complete list of motifs identified only by either XAI or differential analysis.

